# Rearing, dissection, and temporal transcriptomic profiling protocols to study density-dependent phenotypic plasticity in *Schistocerca* (Insecta: Orthoptera)

**DOI:** 10.64898/2026.02.17.705994

**Authors:** Maeva A. Techer, Vivian A. Peralta Santana, Brandon Woo, Richelle Marquess, Christopher Brennan, Audélia M. C. Mechti, Jackson B. Linde, Spencer T. Behmer, Gregory A. Sword, Hojun Song

## Abstract

This protocol generates gregarious and solitarious density-dependent phenotypes in multiple *Schistocerca* species under controlled environmental conditions. It describes cage setup, feeding, animal handling, and sterile dissection workflows to isolate nervous, chemosensory, gut, fat body, and female reproductive tissues from nymphs and adults. It emphasizes rapid tissue stabilization and RNase-control practices for downstream single-tissue DNA and RNA analyses.

**Graphical abstract:** 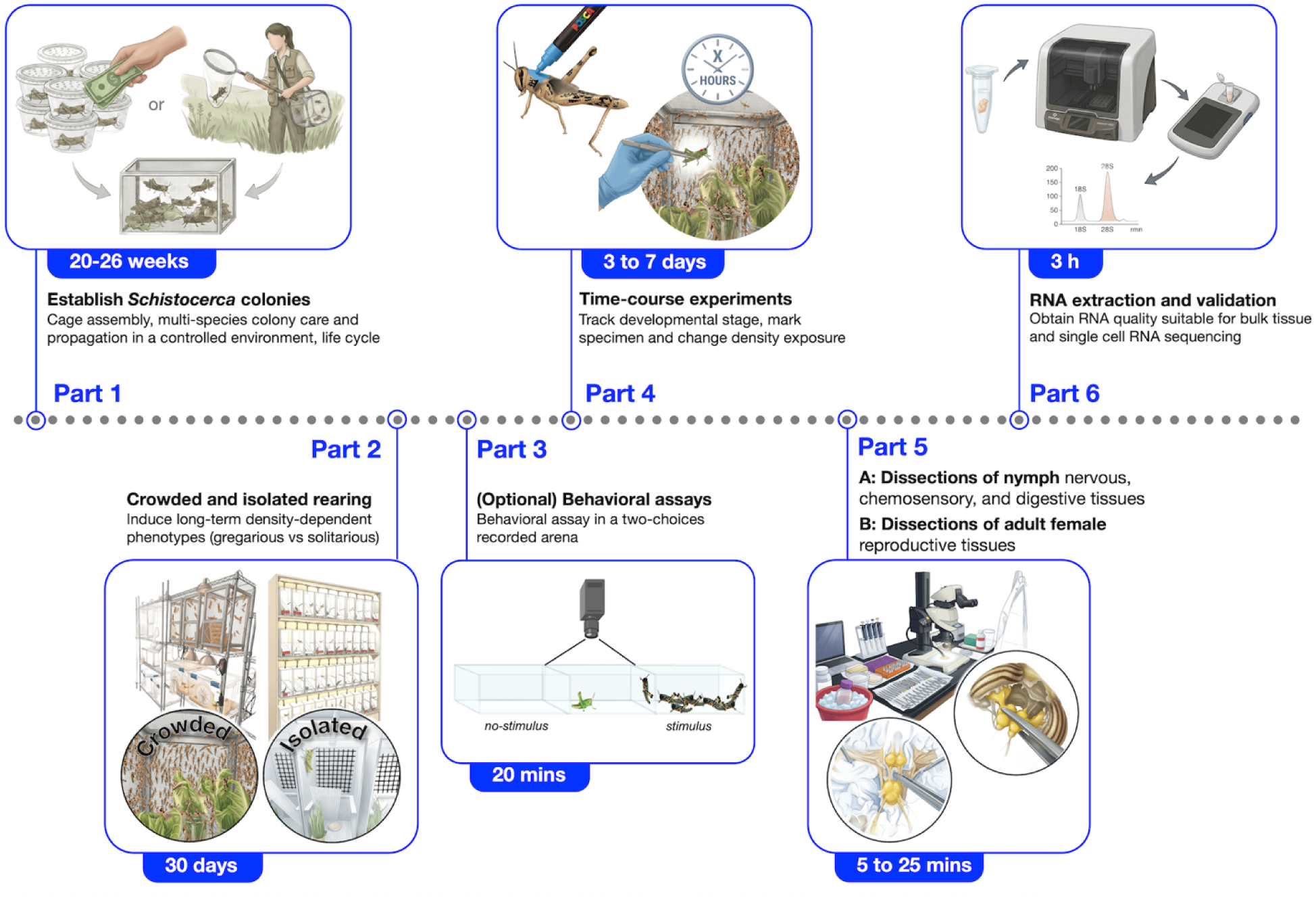

## Before you begin

Locust phase transitions in response to changes in conspecific density represent one of the most impressive phenotypic transformations observed in nature ^1^. Short-term increases in density elicit rapid neural and behavioral responses within hours ^2,3^. Prolonged exposure to tactile, visual, and chemosensory cues leads to profound phenotypic changes in insect morphology, physiology, collective and feeding behavior. Ultimately, these density-driven responses can shift normally sedentary grasshoppers found at low population density (**solitarious phase**) toward collective groups of migrating locusts, manifested as coordinated hopper bands and adult swarms (**gregarious phase**), and this phenomenon is known as locust phase polyphenism. For the past 100 years, since Sir Boris Uvarov proposed the phase theory ^4,5^, our understanding of the proximate mechanisms of locust phase polyphenism has grown tremendously ^6–8^. Much of our current understanding of the physiological, endocrine, genetic, and epigenetic mechanisms underlying locust phase change derives from two established model systems: the Migratory locust (*Locusta migratoria* (Linnaeus)) and the Desert locust (*Schistocerca gregaria* (Forskål)) ^4,9^. Laboratory rearing protocols refined since the 1950s have demonstrated that phase transitions are dynamic and experimentally inducible across all developmental stages ^10,11^. Nevertheless, locusts are not prominently featured in phenotypic plasticity literature ^12^, despite a few previous attempts to highlight locust phase polyphenism as an attractive system ^13^. The main objective of this protocol is to promote locust phase polyphenism as a renewed model system for studying phenotypic plasticity by placing it in the modern theoretical framework of plasticity research.

Compared to other types of polyphenism known in insects ^14,15^, locust phase polyphenism has several attractive features that collectively make this phenomenon ideal for phenotypic plasticity research. First of all, alternative phenotypes are induced by a single environmental cue, local population density, and this has been definitively established for over a century ^16^. Second, density can affect many traits, including behavior, nymphal coloration, morphometric ratios, life history traits, physiology, ecology, biochemistry, and molecular biology, allowing the opportunity to examine the complex nature of phenotypic plasticity. Third, locusts are easy to maintain as laboratory colonies, and the density-dependent alternative phenotypes can be easily induced in the lab at any developmental stage by simply manipulating rearing density. Fourth, the proximate mechanisms of locust phase polyphenism are exceptionally well-characterized in some species, especially in terms of the neurobiological basis of behavioral shift in the Desert locust ^17^, which provides a reference for comparison for other species. Fifth, because locust phase polyphenism has evolved independently in multiple locust species ^18^, the study of locusts naturally lends itself to a comparative framework. Recent studies have shown that closely related locust species vary substantially in both the magnitude and speed of **gregarization** (low-to-high density transition) and **solitarization** (high-to-low density transition) ^18,19^. Moreover, non-swarming grasshopper species closely related to locusts are known to exhibit varying degrees of density-dependent phenotypic plasticity 20, which allowsteasing apart the evolution of individual reaction norms in a phylogenetic framework. Sixth, genomic resources and functional genetics tools are rapidly becoming available for locusts, with the recent release of multiple high-quality genome assemblies (Techer *et al*. in prep), which will help elucidate the molecular basis of phenotypic plasticity. Finally, because locusts are important agricultural pests of biblical proportion, affecting up to one-fifth of the global human population ^20^, understanding the nature of phenotypic plasticity has a direct impact on improving human livelihoods.

That being said, several biological and ecological caveats complicate direct comparisons across species because *Schistocerca* species differ in developmental timing, life history, and climatic adaptation (e.g., presence or absence of diapause), and span habitats from tropical rainforests to dry grasslands and deserts ^21^. Because transcriptional responses to crowding or isolation can occur rapidly and involve multiple small nervous, chemosensory, and reproductive tissues, it is essential to establish methods that precisely control rearing density, track developmental stage, and coordinate behavioral changes with sampling. Despite the widespread use of *S. gregaria*, standardized multi-species protocols implemented under a unified laboratory condition (an artificial common garden) for time-course and tissue-specific experiments applicable to Acrididae remain limited.

### Innovation

Here, we present a comprehensive workflow organized around six sequential steps: (1) establishment of field-collected or laboratory-derived founding *Schistocerca* populations, (2) long-term crowded and isolated rearing, (3) behavioral assays to quantify phase-related responses, (4) time-course experiments during early and late transcriptional response windows, (5) rapid dissection of low-input nervous, chemosensory, and reproductive tissues, and (6) standardized preservation and processing of RNA for transcriptomic analyses. The workflow is designed to be accessible to researchers with limited prior experience in locust and grasshopper rearing while remaining flexible for comparative applications across Acrididae.

This workflow has been developed and validated through extensive multi-species rearing across the genus *Schistocerca*, including three major locust species, the Desert locust (*S. gregaria*), the Central American locust (*S. piceifrons* (Walker)), and the South American locust (*S. cancellata* (Serville)), and three non-swarming grasshoppers (*S. americana* Drury*, S. serialis cubense* (Saussure)*, S. nitens* (Thunberg)). While optimized for comparative temporal transcriptomic profiling, the procedures can be adapted for other molecular applications requiring low-input tissues, not to mention a wide range of behavioral, physiological, or biochemical comparative analyses.

### Institutional permissions

Researchers should ensure compliance with institutional, state, and national regulations governing the transport and maintenance of live insects. In the United States, the interstate or international movement of locusts and grasshoppers is regulated by USDAAPHIS PPQ, and appropriate permits must be obtained prior to initiating this work. *S. gregaria* locusts have been reared in the Texas A&M University Biological Control Quarantine Laboratory (USDA-APHIS-PPQ containment facility #1250) since 2021 and imported under a USDA permit (P526P-19-02151) from the Arizona State University’s Global Locust Initiative (GLI) colony managed by Dr. Arianne Cease. This colony was imported to the U.S. from a long-term colony maintained at the University of Leicester (U.K). The *S. piceifrons* colony was established from an outbreak population in Yucatan, Mexico, imported under a USDA permit (P526P-15-03851), and has been reared in the same facility since 2015. The *S. cancellata* colony was established from an outbreak population in Catamarca province, Argentina, at the Centro de Estudios Parasitológicos y de Vectores (CEPAVE) in La Plata, Argentina. This colony was imported to Texas A&M in 2024 under a USDA permit (P526P-21-06855). The *S. americana* colony was established from a population collected in 2010 in Brooksville, Florida. The *S. serialis cubense* colony was established from a population collected in 2011 in Islamorada, Florida Keys. Finally, the *S. nitens* colony was established from a population collected in 2015 in Terlingua, Texas.

Collection of grasshoppers from public lands, including national, state, or local parks, may require additional permits from the relevant land management agencies. Researchers should verify and obtain all necessary collection permissions before field sampling.

### Preparation of laboratory-crowded and isolated “common garden” rooms

In regions where locust species are not native or established, rearing may be permitted only in a certified quarantine facility. Such facilities may include escape-proof rooms with restricted access, negative air pressure to prevent accidental release, double containment during handling and transport, and strict rules for waste management. Regulatory requirements vary by country and institution and must be confirmed before initiating rearing. The protocols provided here include containment provisions for working under quarantine conditions.

The following protocols work well for species that have been in captivity for successive generations. When working with wild-caught insects, one must carefully consider the potential for pathogens they may carry, which may be transmitted to existing colonies. It is best to quarantine wild-caught insects for at least a full generation before introducing them into the existing colonies.

**Timing:** 2 days (variable depending on cage number, species targeted, and room availability)

**1.** Designate dedicated, climate-controlled rooms for common-garden rearing.

**a.** Assign separate rooms for long-term crowded colonies and for isolated cage-rearing spaces.
**b.** Ensure the rooms can be closed, monitored, and are accessible only by authorized personnel.
**2.** Set standardized environmental conditions appropriate for *Schistocerca* rearing.

**a.** Set the room temperature to a range between 32-34°C and monitor with thermometers.
**b.** Install timers to maintain a consistent 12 h light:12 h dark photoperiod.
**c.** Monitor humidity continuously with hygrometers to be kept stable and below 50%.

**NOTE:** A temperature of 30°C has typically been recommended for general grasshopper and locust rearing ^10^, but our independent observations indicated an absence or delayed molting for last instars when the daily temperature was set lower than 32°C.

**3.** Prepare the **crowded** rearing room to support aggregation and thermoregulation.

**a.** Arrange a heat-source for insects around the cages (either or both):

i. *Light source option 1:* Add one halogen light bulb with a reflector dome above each cage, allowing insects to thermoregulate.
ii. *Light source option 2:* Add LED hanging strips above the cages, arranged so three are available per shelf unit.
iii. Add radiant-heat side panels at the back of the cage to provide a uniform heating source for insects.
**b.** Ensure uniform exposure to light and airflow across cages.

**c.** Leave sufficient space between shelving levels to facilitate cleaning and colony care.
**d.** Leave sufficient space between shelving units when hosting multiple species.
**4.** Prepare the **isolated** rearing room to minimize unintended sensory stimulation.

**a.** The room should be well separated from the crowded room, particularly to avoid exposure to olfactory cues.
**b.** Set up shelves that will support cage spacing for uniform lighting access using LED strip lights.
**c.** Maintain stable, low-disturbance, positive airflow using clean ventilation that feeds into each isolated cage.
**5.** Minimize escape risk in all rearing rooms.

**a.** Seal gaps around doors, vents, and cable entries.
**b.** Place glue traps near room entrances and shelving units where cages will be positioned.
**c.** Ensure doors remain closed when rooms are unoccupied.
**d.** Have small containers (e.g., zip-lock bags), insect nets, and a vacuum readily available to capture and immediately freeze any escapees at -20 °C.
**6.** Arrange a waste management system with a freezer and an autoclave in an easily accessible location.

**a.** Designate a -30 °C freezer for temporary storage of biological waste (e.g., deceased insects) before disposal.
**b.** Ensure access to an autoclave for the sterilization of reusable materials and decontamination of waste before removal from the facility.
**c.** Establish and follow a standard operating procedure (SOP) for the rearing facility, approved by local or federal agencies.
**7.** Establish room-level containment and hygiene practices.

**a.** Place clear signage indicating live insect work and regulatory compliance as required.
**b.** Keep cleaning supplies (e.g., bleach, ethanol, vacuum, wipes) available within each room.
**d.** Designate lab coats or protective clothing for use exclusively within rearing rooms.
**e.** Implement a color-coding and tracking system with labels for multi-species work.
**8.** Once environmental conditions are stable for 24h, the room is ready for cage installation and insect introduction.

### Preparation of locust and grasshopper cages

The cage format recommendations are based on >15 years of successful long-term rearing and experimental work on *S. gregaria* (two strains), *S. piceifrons*, *S. cancellata*, *S. americana*, *S. serialis cubens*e, and *S. nitens* under crowded conditions. Gregarious locusts rapidly chew through organic materials, rendering many soft or fabric insect cages unsuitable.

**Timing:** 10-30 min (depending on the cage type and custom-designed pre-assembly status)

**1.** Select cage type based on developmental stage and density condition (Fig. 1A).

**a.** For **long-term crowded** 1st to last instar:

**i.** Use cubic aluminum-frame mesh cages (12″ × 12″ × 12″) for moderately crowded rearing or fine life-cycle tracking. 3-4 egg cups can be placed inside at once (Fig. 1B).
**ii.** Use weather-tight clear plastic boxes (24” x 18” x 15 ¼”, ∼80 qt) for highly crowded rearing. 7-8 egg cups to be placed inside at once (Fig. 1B).
**iii.** Use fabric mesh cages only for sedentary or small-bodied species (e.g.,
*S. nitens*) when chewing damage is minimal.
**b.** For **long-term crowded** last instar and adults:

**i.** Use tall aluminum-frame cages (12” x 16” x 24”) with 20 × 20 aluminum mesh and a transparent Plexiglass access door (Fig. 1B).
**ii.** Prepare an elevated cage bottom with 0.4″ mesh openings positioned above a removable tray to allow frass to fall while preventing escape.

**Figure 1:**
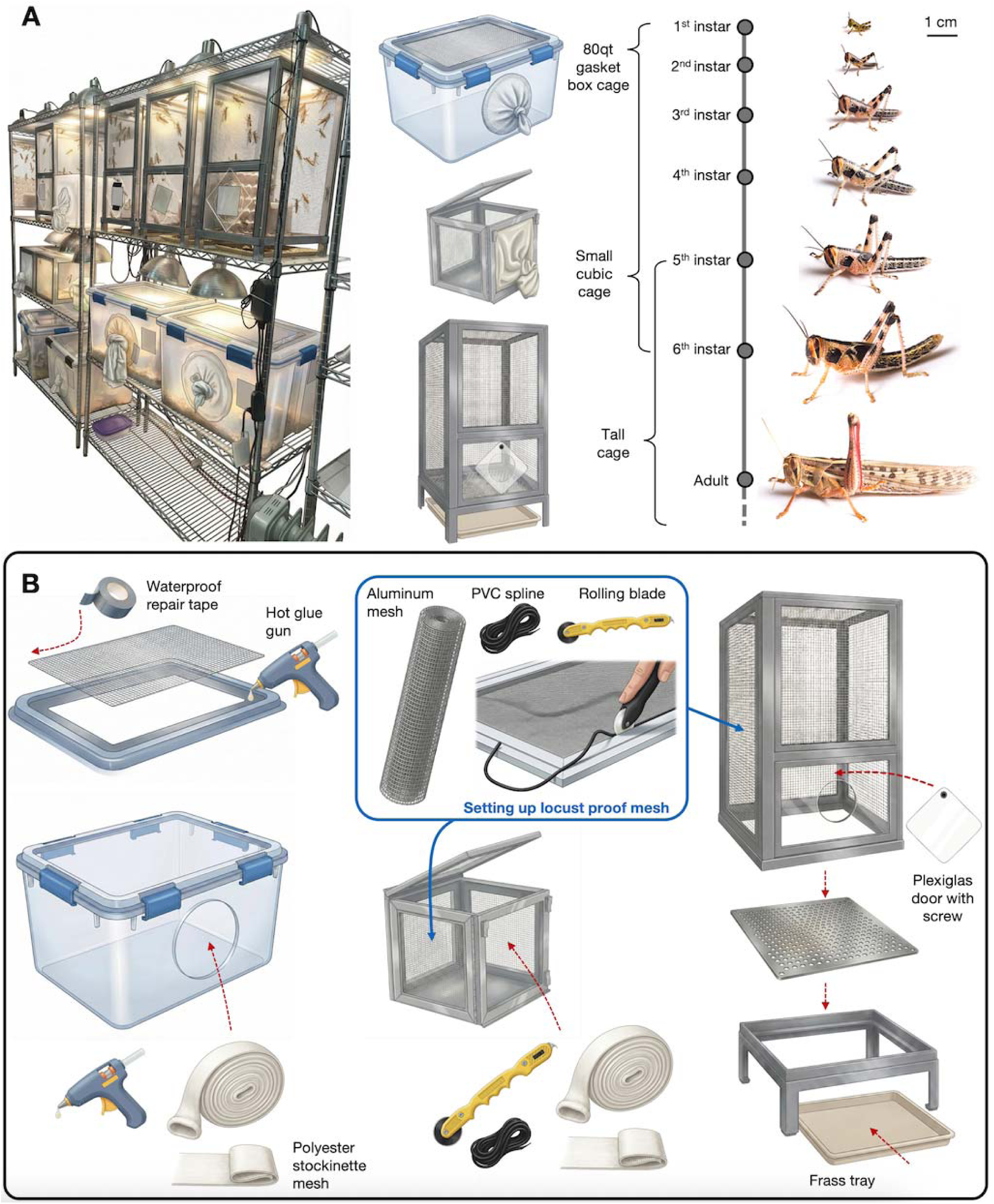
Cage systems for crowded rearing of *Schistocerca* nymphs and adults. (A) Example of shelving organization in a common-garden rearing room at Texas A&M University, Locust Quarantine Facility, with one shelf per species and cage types selected according to developmental stage, with light bulbs above or LED band lights. Hatchlings and nymphs (1st to last instar) are maintained in vented clear gasket boxes or cubic aluminum-frame mesh cages, whereas adults are housed in taller cages that allow vertical movement, wing expansion, and oviposition. (B) Overview of cage assembly principles for high-density locust and grasshopper rearing. Commercial cage formats are custom-modified to improve containment, facilitate daily maintenance, and allow efficient cleaning during long-term colony propagation.

**NOTE:** An alternative is to keep a single large cage type for an entire cohort (e.g., from 1st instar to adults), and such a setup has been adopted in various labs, including the Arizona State University GLI lab.

**c.** For **long-term isolation** of nymphs and adults:

**i.** Use a custom-designed rigid plastic cage (10.16 × 10.16 × 25.4 cm) with a transparent front door and a 0.5-inch diameter airflow port at the top (Fig. 2).

**Figure 2:**
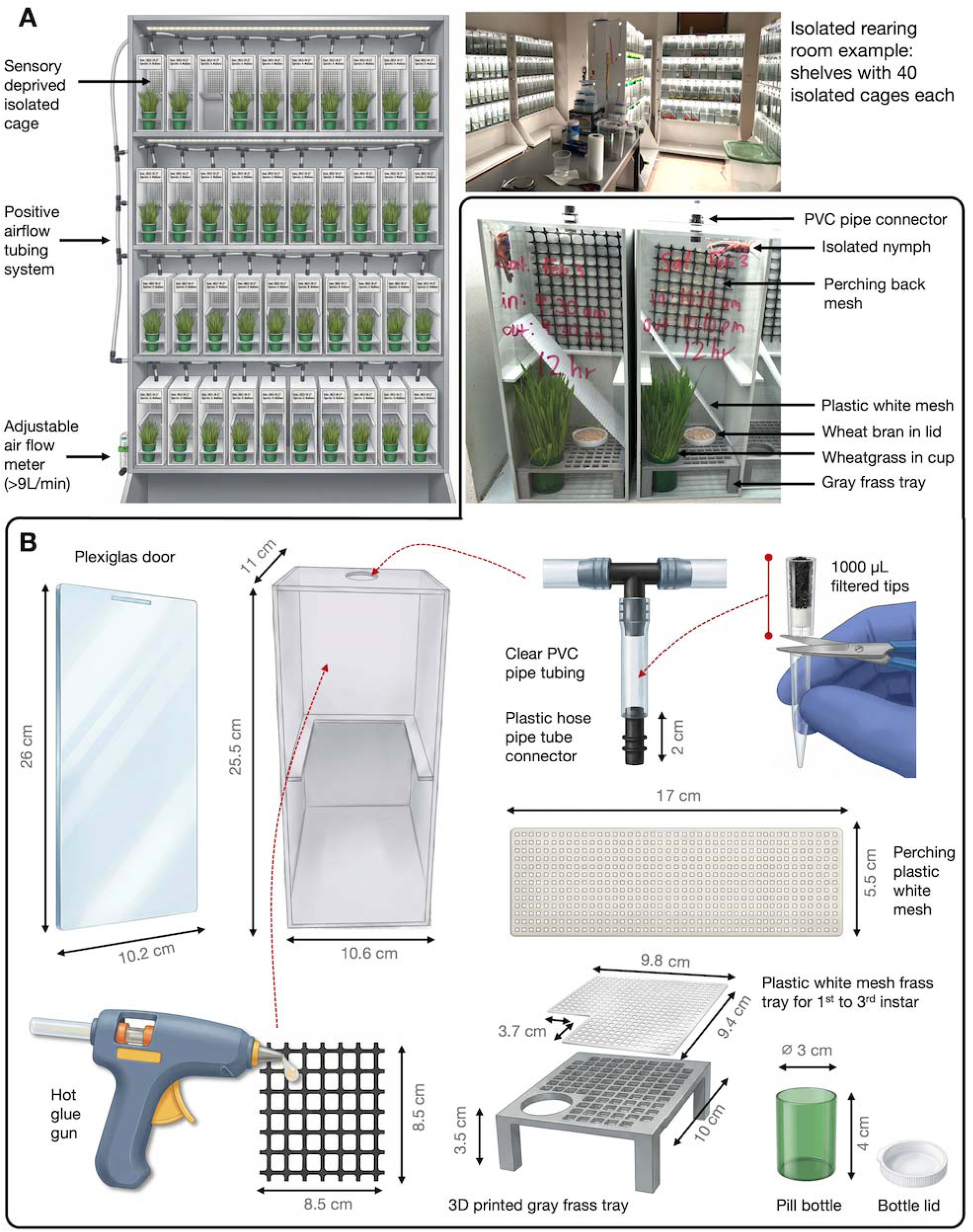
Cage systems for isolated rearing of *Schistocerca* nymphs and adults. (A) Custom-designed individual cages are used for long-term isolated rearing under common-garden conditions. Each cage houses a single nymph or adult and is arranged on dedicated shelving to ensure complete visual, tactile, and olfactory isolation from conspecifics. (B) Key structural features of isolated cages include controlled positive ventilation, filtered ventilation, internal perching surfaces, and removable components that facilitate daily care, feces removal, and experimental handling while minimizing unintended sensory stimulation.

**NOTE**: Custom-designed and/or 3D-printed components should be ordered and prepared in advance.

**2.** Restrict use of temporary cages to transport or short-term assays only:

**a.** Use heavyweight polypropylene deli containers (16 oz or 32 oz) with pierced LDPE lids (<0.1″ holes).
**b.** Use Port-A-Bug containers or 4–8 qt clear storage round storage containers as needed.
**c.** Always place temporary cages inside a secondary container to ensure double containment during transport.
**3.** Assemble sides, panels, and ventilation surfaces for **crowded** cages.

**a.** Install aluminum screen mesh (∼20 × 20) across cage openings using spline, hot glue, or equivalent fasteners appropriate to the cage design (Fig. 1B).
**b.** Install a double-layer knitted polyester stockinette access sleeve for feeding and maintenance.

**i.** Cut a sleeve opening (as required by cage type) and attach two layers of stockinette securely from the inside of the cage (using spline or hot glue).
**ii.** Ensure the sleeve is long enough to be knotted when not in use and to allow feeding and cleaning without creating a direct escape path.
**c.** Reinforce all mesh seams and edges with waterproof repair tape (or equivalent) to eliminate gaps (Fig. 1B).
**d.** Inspect corners, hinges, and lid interfaces for openings and seal as needed.
**4.** Add internal structures and waste management components for **crowded** cages (Fig. 3).

**a.** Add egg carton perches to nymph and adult crowded cages to increase climbing surface and provide protection during molting.
**b.** For nymph-crowded cages, aspen wood shavings can be added as substrate to absorb moisture and reduce mold.
**c.** For tall adult crowded cages, place a removable tray beneath the cage to collect frass and debris (Fig. 1B).
**5.** Assemble ventilation and internal structures for **isolated** cages (Fig. 2B).

**a.** Cut clear PVC tubing to the appropriate length and insert it into the top airflow port to ensure stable, low-disturbance ventilation.
**b.** Seal the external end of the tubing with either a knitted stockinette sleeve or a 1,000 µL filtered pipette tip containing activated charcoal.
**c.** Glue a square of black vinyl-coated hardware cloth with ½ in mesh (8.5 x 8.5 cm) to one inner wall of the cage to provide vertical perching.
**d.** Insert a white plastic mesh plate (approximately 5 cm × 10 cm) to provide a perching and resting surface.
**e.** Place a custom 3D-printed gray tray table with a central opening at the cage bottom to support food cups and allow frass to fall away from the insect.
**f.** For the 1st to 3rd instar, a white plastic canvas mesh piece can be cut as shown in Fig. 2B to prevent insects from interacting with the frass collection section of the cage.
**6.** Finalize containment checks and labeling.

**a.** Confirm that lids latch tightly and that sleeves, doors, and closures return to a sealed position after handling.
**b.** Repair or replace any mesh with holes, loose tape, or compromised seams immediately.
**c.** Label each cage with species, density condition, colony or experiment ID, and date using color-coded tape for multi-species work.
**7.** Clean and decontaminate cages before reuse. Following insect transfer or at the end of cage use, all cages must be fully cleaned and dried before reassembly.

**a.** Remove all stockinette sleeves and mesh components; discard or wash thoroughly if reusable.
**b.** Discard all soiled organic internal components, including egg cartons, substrates, and organic debris (Fig. 3).
**c.** Submerge the entire cage, cups, and 3D printed tables in 3% bleach solution or Rescue™ disinfectant for 30 min.
**d.** Scrub all surfaces with a brush to remove frass, residue, and remaining debris.
**e.** Rinse thoroughly with clean water to remove all disinfectant residues.
**f.** Allow cages, cups, and 3D parts to dry completely before reassembling or replacing missing components.

**Figure 3:**
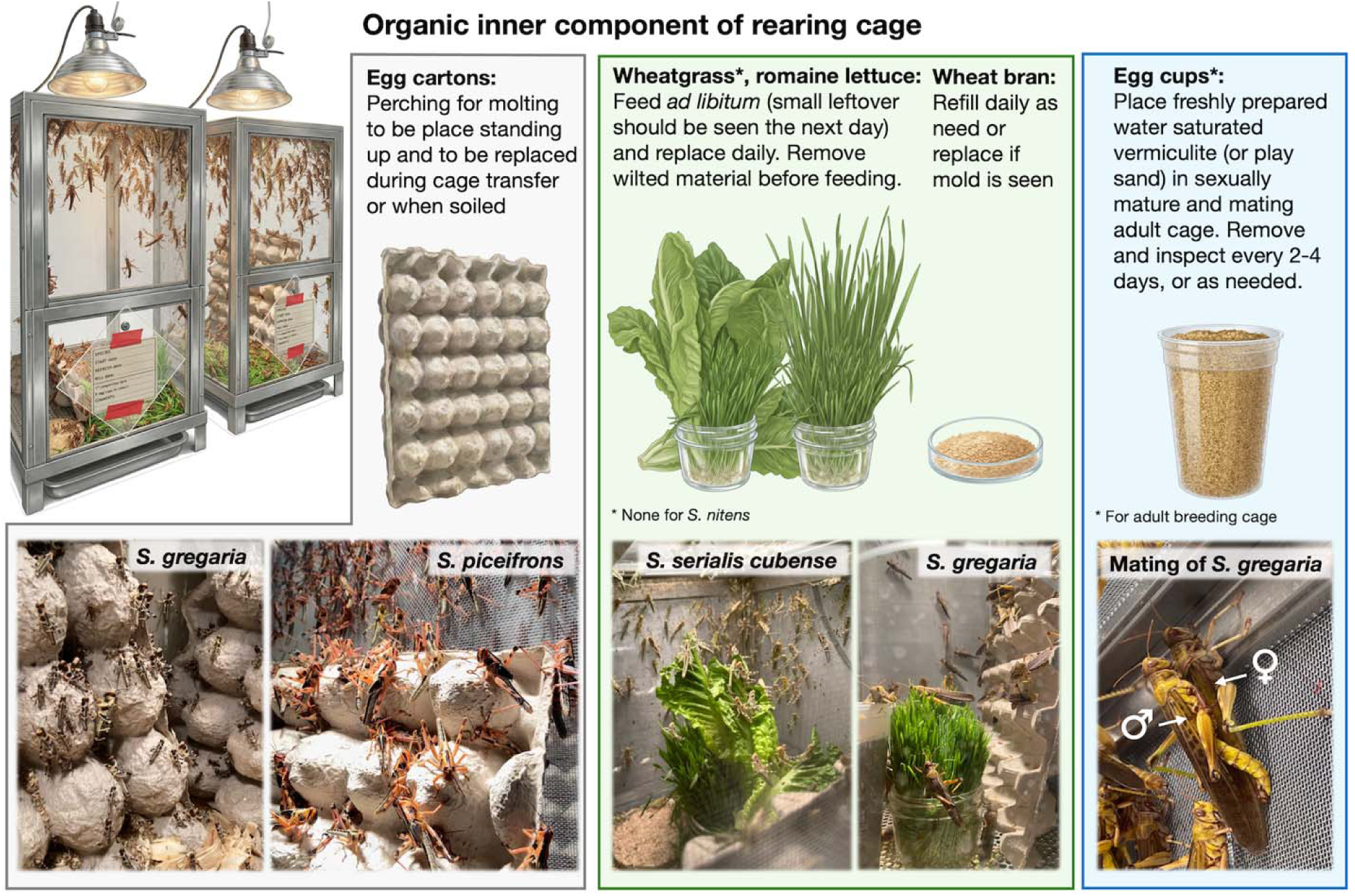
Internal organic cage components used in crowded rearing for multiple *Schistocerca* species. Food sources are identical to those used in isolated cages but are given in size- and layout-adapted containers to accommodate spatial constraints. Make sure to use organic romaine lettuce (or any other produce) to avoid potential pesticide residues that may be present in store-bought produce.

### Preparation of sterile locust saline solution

**Timing:** 30 min (plus autoclaving time)

**1.** Add 900 mL of Milli-Q water to a sterile 1 L autoclavable glass bottle.
**2.** Add the reagents listed in Table 1 to the bottle.

**a.** Weigh each reagent accurately using an analytical balance.
**b.** Add reagents sequentially while stirring to prevent precipitation.
**3.** Stir the solution continuously using a magnetic stir bar until all salts and sugars are fully dissolved and no visible particles remain.
**4.** Measure the pH of the solution using a calibrated pH meter or pH indicator strips.

**a.** Adjust the pH to 7.4 by adding hydrochloric acid (HCl) or sodium hydroxide (NaOH) dropwise while stirring.

**Table 1.**
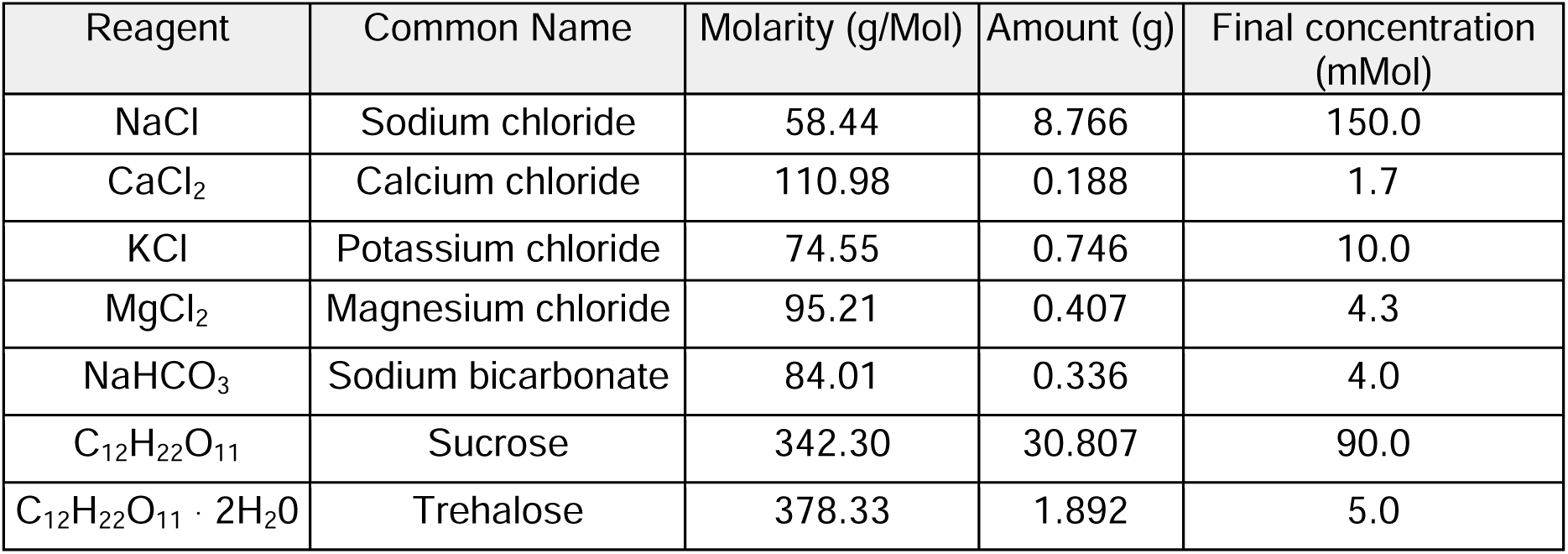
Reagents used to make 1L of locust saline solution.

**CRITICAL:** Add acid or base slowly and mix thoroughly after each addition to avoid overshooting the target pH.

**5.** Bring the final volume to 1 L by adding Milli-Q water.
**6.** Autoclave the solution using a standard liquid cycle.
**7.** Allow the solution to cool to room temperature and store at 4 °C.

**Pause point:** The sterile locust saline solution should be used within 3-5 days. Because the solution contains sugars, extended storage may increase the risk of microbial contamination.

**Note:** Before use, visually inspect the solution for turbidity or particulate matter and discard if contamination is suspected.

### Preparation of the dissection bench

**Timing:** 35 min (plus overnight cooling of locust saline solution and dissection plates)

**1.** Set up a stereo microscope and appropriate lighting on a clean, flat surface (bench or table). Position the microscope centrally on the bench to allow comfortable access during dissection.
**2.** Set up a microbead sterilizer and turn it on 30 min before the start of a dissection.
**3.** Prepare all dissection tools and fine supplies in advance and autoclave them before use (see Figure 4 for an example of the dissection setup).
**4.** Clean the dissection station and stereo microscope platform with 3% bleach solution, then rinse thoroughly with distilled water.

**a.** Dry all surfaces with clean paper towels.
**b.** Repeat the rinse step to remove residual bleach.

**Figure 4:**
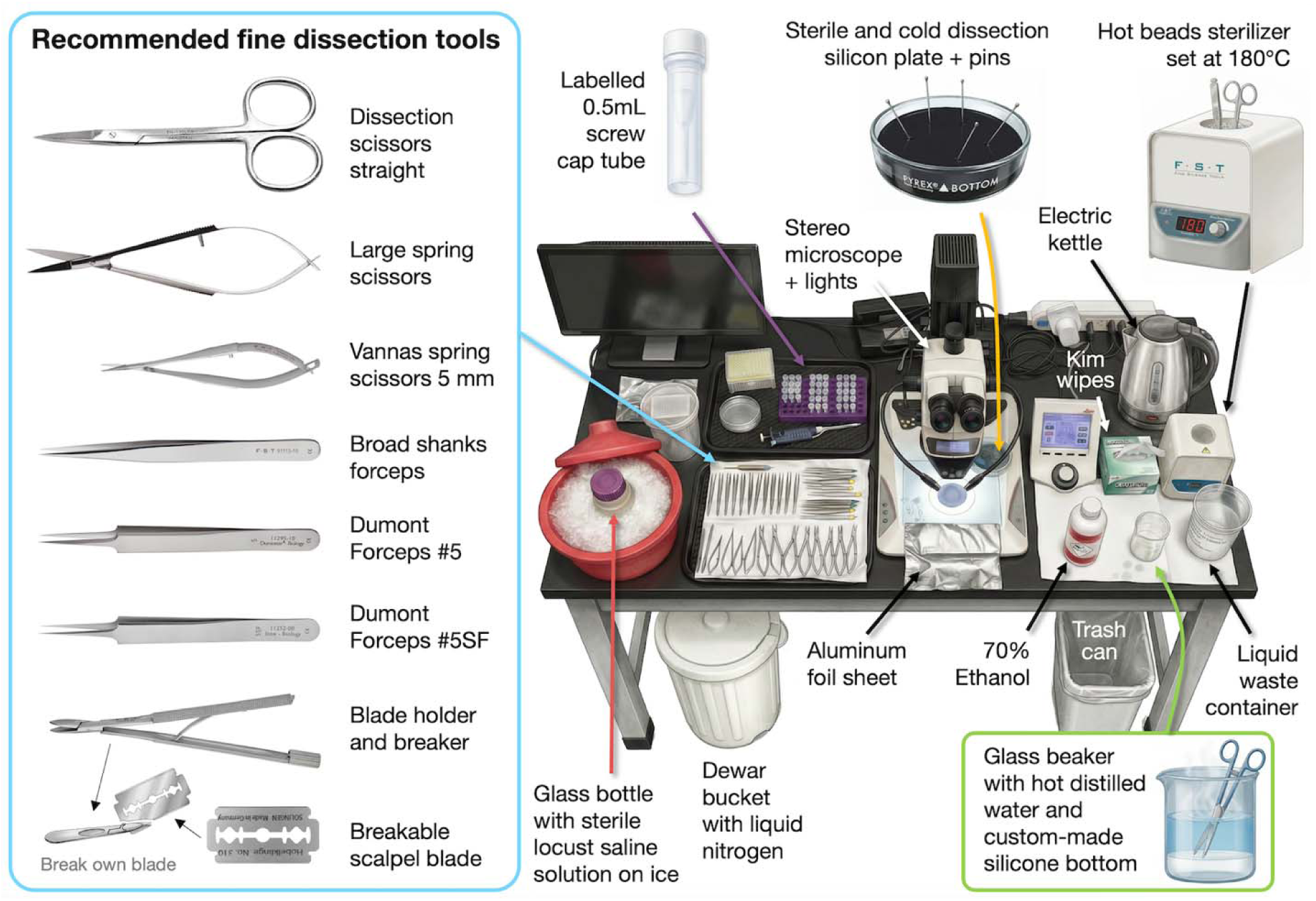
Bird’s-eye-view of a dissection bench setup for rapid and successive tissue isolation from live locust and grasshopper specimens. The arrangement highlights recommended fine dissection tools used in Parts 5A and 5B, which should be prepared in multiple sterile sets depending on the time available for cleaning/sterilization, tissues targeted, and the number of consecutive dissections performed.

**CRITICAL:** If tissues will be used for RNA analysis, wipe the dissection station and all tools with RNase decontamination solution (e.g., RNaseZap) after cleaning and immediately before starting the dissection.

**5.** Arrange fine forceps, spring scissors, insect pins, and other dissection tools within easy reach of the dominant hand (Fig. 4).
**6.** Place an ice bucket, sterile 0.5mL screwcap standing microcentrifuge tubes, and a waste container nearby to minimize movement during dissection.
**7.** Label screw-cap tubes and place them upright on a chilled tray. Assign one tube per tissue per individual specimen.
**8.** Fill a heat-resistant, silicone-bottomed instrument beaker with hot distilled water and place it on the dissection bench for tool cleaning purposes (Fig. 4).
**9.** Prepare a liquid nitrogen container with long tweezers so that labeled tubes can be rapidly submerged immediately after tissue removal.
**10.** Place the locust saline solution on ice.

**a.** Plan to use approximately 100–200 mL of saline per dissection session.
**11.** Cut a square of aluminum foil approximately 15 × 15 cm and place it near the dissection area.
**12.** Retrieve clean, autoclaved dissection plates as needed (Fig. 4).

**a.** Chill plates at 4 °C for at least 5 h or at −20 °C for at least 2 h before dissection.
**b.** Insert insect pins vertically into the silicone base of one dissection plate to facilitate rapid pinning of specimens.
**13. Optional**: Use a timer when beginning the dissection to monitor total handling time and minimize RNA degradation associated with prolonged tissue exposure.

**Note:** This protocol requires handling with clean gloves. If gloves become contaminated before or during dissection (e.g., contact with regurgitate or frass), either wipe gloves thoroughly with 70% ethanol or replace them immediately. To reduce contamination when working with low-input tissue, at a minimum, wear a lab coat and face mask, and tie long hair back.

## Key resources table

1 Key resources table
| REAGENT or RESOURCE | SOURCE | IDENTIFIER |
| --- | --- | --- |
| Antibodies |  |  |
| Bacterial and virus strains |  |  |
| Biological samples |  |  |
| Chemicals, peptides, and recombinant proteins |  |  |
| Ethanol, Absolute (200 Proof), Molecular Biology Grade | ThermoFisher | BP2818-100 |
| Rescue pet solution | Amazon | B00B9G3ZJM |
| ddH2O or autoclaved water | Texas A&M University |  |
| NaCl | Sigma-Aldrich, USA | S3014-500G |
| CaCl<sub>2</sub> (Sigma-Aldrich, USA, Cat. No.:) | Sigma-Aldrich, USA | C1016-500G |
| KCl (Sigma-Aldrich, USA, Cat. No.:) | Sigma-Aldrich, USA | 746436-500G |
| MgCl<sub>2</sub> (Sigma-Aldrich, USA, Cat. No.:) | Sigma-Aldrich, USA | M4880-100G |
| NaHCO<sub>3</sub> (Sigma-Aldrich, USA, Cat. No.:) | Sigma-Aldrich, USA | S5761-500G |
| Sucrose (Sigma-Aldrich, USA, Cat. No.:) | Sigma-Aldrich, USA | S0389-500G |
| Trehalose (Sigma-Aldrich, USA, Cat. No.:) | Sigma-Aldrich, USA | T9531-10G |
| RNAseZap™ RNase Decontamination Solution | Invitrogen, USA | AM9780 |
| Critical commercial assays |  |  |
| Maxwell® RSC simplyRNA Tissue Kit | Promega | AS1340 |
| High Sensitivity RNA ScreenTape | Agilent | 5067-5579 |
| High Sensitivity RNA ScreenTape Ladder | Agilent | 5067-5581 |
| High Sensitivity RNA ScreenTape Sample Buffer | Agilent | 5067-5580 |
| Qubit™ RNA HS Assay Kit | Invitrogen | Q32852, Q32855 |
| Deposited data |  |  |
| Experimental models: Cell lines |  |  |
| Experimental models: Organisms/strains |  |  |
| Oligonucleotides |  |  |
| Recombinant DNA |  |  |
| Software and algorithms |  |  |
| Other |  |  |
| Climatic control incubator chamber | Percival Scientific | I-41NL |
| Multiple Shared Use Autoclave | TOMY | SX-700/500/300 |
| -30°C Freezer (any brand) | FisherScientific |  |
| 4°C Freezer (any brand) | Lowes |  |
| ddH2O water dispenser | Texas A&M University |  |
| Stainless steel shelves (48 x 24 x 86 inch) | Uline | H-2946-86 |
| Custom-designed tall rectangular cages (30.5 x 30.5 x 61 cm) with an aluminum frame, plexiglass door, and elevated sieve bottom | Inquire engineer department or a private business |  |
| Clear plastic storage containers (61 x 45.7 x 38.1 cm) with tight-fitting latches | Sterilite, USA | 19384304 |
| Cubic collapsible cages (30.5 x 30.5 x 30.5 cm) with aluminum frame | Bioquip, USA | 1450BC |
| Polyester waffle mesh stockinette 4" (any brand) | Kinship Comfort Brands | B07DYVXWRF |
| Indoor/Outdoor PAR38 flood halogen light bulb 90W | Philips | 419382 |
| 10 or 8.5 in. dome light reflector | Home depot | UR13 Y1T |
| Boostgro LED Seed Starter Grow Lights (3ft) Full Spectrum (5000k Daylight + 660nm Red + 450nm Blue. | DYMOND | 850061919168 |
| Radiant Heater - side mount 11 x 40 | Sweeter Heater | 4G-NEI8-10XI |
| Aluminum insect screen | Home depot | SKU 1001541819 |
| Black window spline (4.1mm) | Home depot | SKU 776262 |
| Screening roller knife | Screen Tight, USA | ROLLERKNIFE |
| Rectangular cafeteria tray 10" x 14" (any brand) | Carlisle FoodService | B003SSR8C4 |
| Handheld vacuum | Walmart | 3106701139 |
| Paper egg cartons (11.75"L x 11.75"W x 5.5"H) (any brand) | MT Products Store | UPC 646847486524 |
| PYREX Reusable Petri dishes | Thermo Fisher Scientific, USA | 08-747D |
| Colored masking tape | Walmart |  |
| Thermometer and hygrometer (smart) | Govee | B0872ZWV8X |
| Uni Posca acrylic water-based paint marker, medium point | Mitsubishi pencil, Japan | PC-5M15C |
| Large glue traps | Walmart | B0757F2D74 |
| 8 Qt. Clear Round Polycarbonate Food Storage | Cambro | 214RFSCW8 |
| Container |  |  |
| 32oz Heavy duty Deli cups | TSK supply |  |
| Vented lid for deli cups and insects | TSK supply |  |
| Vermiculite | Uline |  |
| Custom-designed isolated cages (25.5 x 11 x 10.6 cm) | Inquire engineer department or a private business |  |
| Custom-designed isolated shelves (76 x 5.5 x 48 cm) | JAM Designs |  |
| Green plastic cups with lids (pill bottle) | The vial company |  |
| White plastic canvas sheets mesh (any brand) | YOUNTHYE Store | B08C2957RR |
| Black vinyl coated hardware cloth with ½ in mesh | Home depot | 334634505 |
| PVC clear tubing X 0.5" | Home depot | 702361 |
| T-shape joint hose connector | Home depot | 206944256 |
| Filtered pipette tips 1000 µL | VWR | 02-707-404 |
| Premium activated carbon | Marineland | B0002566WY |
| Air flowmeter 1-10LPM with control valve for oxygen air | Hilitand, USA | LZQ-7 1-10LPM |
| LED strip white lights (for isolated shelves) | RadiantFlux Store | B0C96KQ9VQ |
| Organic Romaine lettuce | HEB |  |
| Wheatgrass seeds | Local co-op |  |
| Black germination trays | Greenhouse megastore |  |
| 4-quart colander | Walmart |  |
| Disposable hand sports towel (19.7"L x 27.56" W) (any brand) | Biubumtu | B0D8V6NGKZ |
| Wheat bran flakes | Bulkfoods.com |  |
| Binocular Macroscope | Leica | M205-A |
| Dissection Forceps with broad shanks | Fine Science Tools, USA | 91113-10 |
| Fine Forceps Dumont #5 | Fine Science Tools, USA | 11295-10 |
| Super Fine Forceps Dumont #5SF | Fine Science Tools, USA | 11252-00 |
| Dissection Scissors Straight | Fine Science Tools, USA | 14060-11 |
| Microdissection Vannas spring scissors 5 mm | Fine Science Tools, USA | 91500-09 |
| Blade holder and breaker | Fine Science Tools, USA | 10053-09 |
| Breakable scalpel blades | Fine Science Tools, USA | 10050-00 |
| Plastic sterilization containers | Fine Science Tools, USA | 20840-03 |
| Dissection dishes, large 93 mm diameter, black | Living Systems Instrumentations | DD-90 |
| insect pins/needles, no head #3 | ecologysupplies.com |  |
| 10-100 µL filtered pipettes | VWR |  |
| Microcentrifuge tubes, 0.5mL free standing screw cap | Fisherbrand | 02-681-370 |
| 24-well cell plates | ThermoFisher | 144530 |
| 1L graduated glass bottle | PYREX® | 1396-1L |
| 1L Serialized Class A Cylinders | FisherScientific | 0973030 |
| Instrument beaker with silicon bottom | Fine Science Tools, USA | 00081-00 |
| Kim wipes | Kimberly-Clark Professional™, USA | 06-666 |
| Aluminum foil | Walmart | 11027095 |
| Paper towels | Walmart | 3945859864 |
| Disinfecting Bleach 7.5% | Walmart | 629539844 |
| Magnetic stirrer | FisherScientific | 11676267 |
| Hot Bead Sterilizers | Fine Science Tools, USA | 18000-45 |
| Liquid nitrogen dewar flask | Nalgene®, VWR | 63380-052 |
| Electric water kettle | Walmart | 5329046667 |

### Step-by-step method details

## Part 1: Establish *Schistocerca* colonies

**Timing:** 20 to 26 weeks (variable depending on species, source population, and associated permit)

This step describes how to initiate, propagate, and stabilize *Schistocerca* colonies under controlled laboratory conditions before experimental density manipulation. It ensures synchronized generations, minimizes the introduction of pathogens, and enables accurate life cycle tracking across species.

**1. Identify the source of the founding population.**

a. From established laboratory or commercial suppliers:

i. When egg pods are available, use them directly to initiate the F0 generation and proceed to common-garden rearing.
ii. When egg pods are unavailable, collect or obtain 100–200 adults or late-instar nymphs to establish a breeding population.

**NOTE:** The six *Schistocerca* species described in this protocol do not exhibit egg diapause, allowing continuous breeding under laboratory conditions; however, other univoltine grasshopper species may exhibit egg diapause, which is known to be difficult to break. The wild-caught *S. piceifrons* did exhibit a prolonged (3-month) reproductive diapause as adults, but subsequent generations did not exhibit any reproductive diapause, suggesting that it is feasible to establish this species as a continuously breeding colony. From field collection:

iii. Identify collection sites and seasons in consultation with grasshopper biologists or agricultural extension agents.
iv. Use biodiversity databases (e.g., GBIF, iNaturalist, Orthoptera Species File) to confirm recent species occurrences at the collection site.
v. Confirm species identity with taxonomists.
vi. Sex individuals by examining the genitalia (Fig. 5).

**Figure 5:**
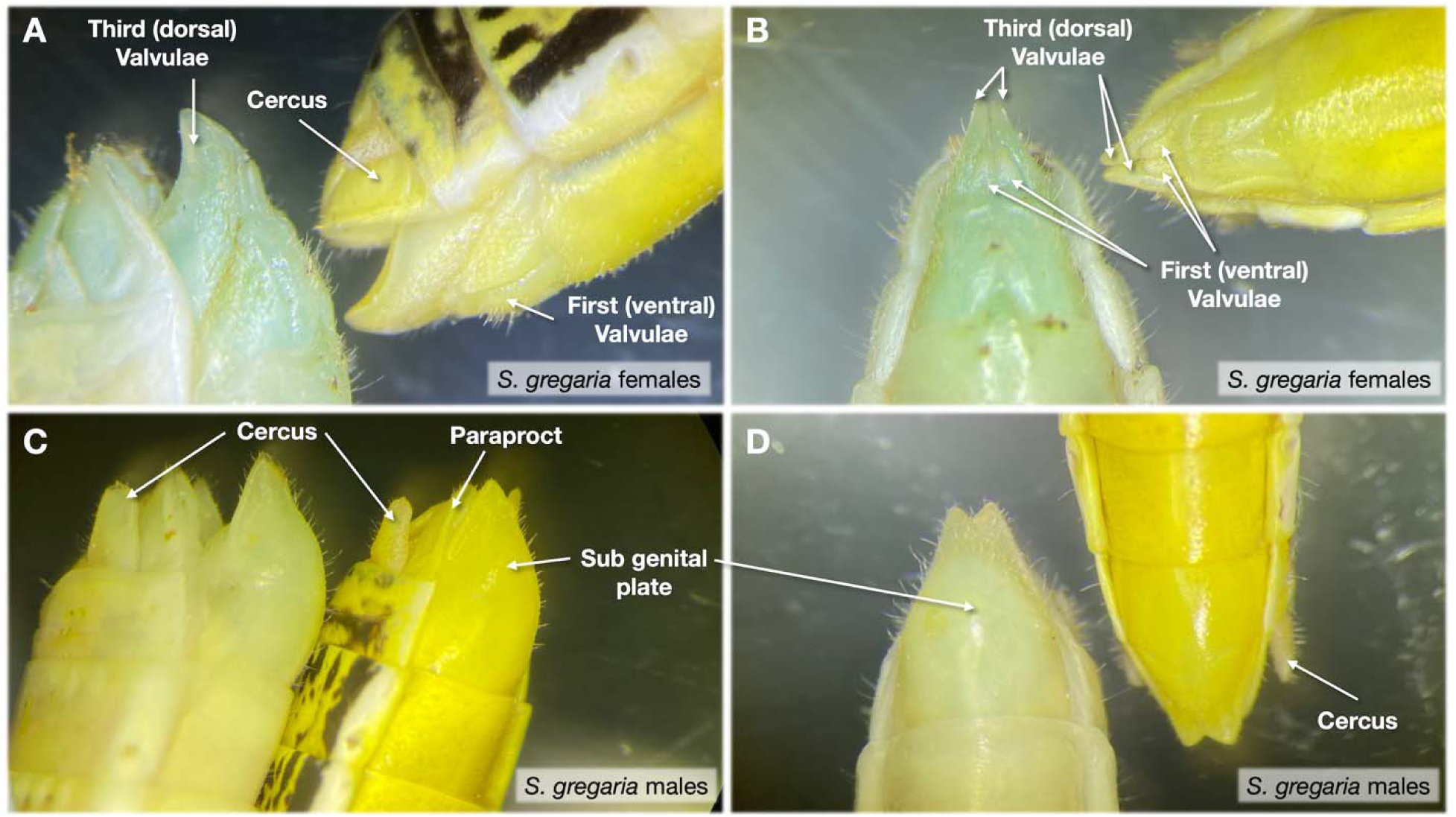
Guide for sex identification and phase differentiation using external genital morphology of last-instar *S. gregaria* nymphs. Female nymphs exhibit a complex ovipositor apparatus composed of multiple valvulae, shown in left lateral (A) and ventral (B) views, in both solitarious (green on the left) and gregarious (yellow on the right) phases. Male nymphs display a smooth, continuous subgenital plate, shown in left lateral (C) and ventral (D) views.

**NOTE:** All *Schistocerca* colonies maintained at Texas A&M University were originally established from field-collected individuals, with the exception of *S. gregaria,* which originated from a laboratory line from the University of Leicester, UK.

**2. Secure permits and determine quarantine requirements.**

a. Obtain all required institutional, state, and national permits for the collection, transport, and rearing of live insects before initiating colonies.
b. When colonies are initiated from laboratory egg pods, transfer egg pods directly into common-garden rearing rooms under standardized conditions.
c. When colonies are initiated from live adults or nymphs, place individuals in temporary quarantine cages (not in common garden space) until laboratory-laid egg pods are obtained.

**CRITICAL:** Permit approval may take weeks to months and should be initiated as early as possible.

**3. Maintain founding insects and monitor colony health.**

a. Monitor colony health and cage integrity.

i. Inspect cages daily for escape risk, abnormal mortality, mold growth, or pest contamination.
ii. Using gloves, count and place dead adults in zip-lock bags, and freeze them at -20 °C overnight.
iii. Spray 70% ethanol on the gloves between cages of the same species.
iv. Change gloves between species cages.
v. For suspicious death rates, check and research any pathogens associated with symptoms of bacterial, fungal, and viral contamination.
b. Supply fresh plants daily or at species- and stage-appropriate intervals. **NOTE:** *S. gregaria, S. piceifrons, S. cancellata, S. americana,* and *S. serialis cubense* thrive with a combination of freshly cut wheatgrass and organic romaine lettuce. Organic cabbage, oatgrass, and sorghum are also suitable sources of fresh food. However, *S. nitens* does not prefer wheatgrass and only feeds on herbaceous plants, such as organic romaine lettuce. All species should be supplemented with wheat bran in a clean petri dish.

i. Remove wilted, contaminated, or uneaten plant material promptly.
ii. Introduce food through stockinette sleeves to minimize the risk of escapees.
c. Maintain cage hygiene

i. For nymph cages: Replace heavily soiled internal structures (e.g., egg cartons, perches, substrate).
ii. For adult cages: throw out the feces that have fallen in the tray and replace the heavily soiled internal structure.
**4. Induce oviposition and collect egg pods.**

a. Monitor adult cages for mating and oviposition activity, including abdominal expansion, probing behavior, or dried egg-foam deposits (Fig. 3 and 6). Sexually mature adult males become yellow in *S. gregaria* (most pronounced), *S. piceifrons* (somewhat), and *S. americana* (somewhat). *S. cancellata* males stridulate with wings when sexually mature.
b. Prepare oviposition cups by filling clear plastic deli cups with fine-grade vermiculite hydrated with distilled water until uniformly moist but not soaked.
c. Place oviposition cups into adult cages and collect them every 3–5 days, depending on cage density and oviposition activity.
d. Remove surface fecal material and inspect cups for egg pods by shallow excavation when necessary or confirm presence with side visibility (Fig. 6).
e. Cover exposed foam plugs lightly with fresh vermiculite and add water if cups have dried much.
f. Label each cup with species name, collection date, and colony identifier. **NOTE:** Egg laying often occurs during the evenings. Additionally, frass can rapidly accumulate at the surface of the vermiculite and needs to be cleaned or replaced to provide fresh laying material to insects.
**5. Incubate egg pods and monitor hatching.**

a. Incubate egg cups in a temperature range between 32-34°C (Fig. 6). As eggs are underground, the need for a light stimulus as a hatching cue is still debated ^22,23^.
b. Check cup moisture daily and add water as needed.
c. Record hatching time according to species-specific developmental windows. Use observations of lab colonies at Texas A&M University’s Locust Quarantine Facility, in Figure 7A, presenting the time for first emergence for four *Schistocerca* species.
**6. Transfer hatchlings and propagate colonies**

a. Transfer hatchlings to nymph cages as soon as possible, providing fresh food and perching/hiding substrate to reduce mortality.
b. Place unopened egg cups containing hatchlings into prepared, latched nymph cages.
c. Open the cups inside the cage and remove the cup lid after shaking it several times to dislodge any remaining nymphs, while maintaining the stocking tight to your arm (Fig. 6).
d. Leave cups in cages for up to 48 h to allow late-hatching nymphs to emerge, then remove them.

**Figure 6:**
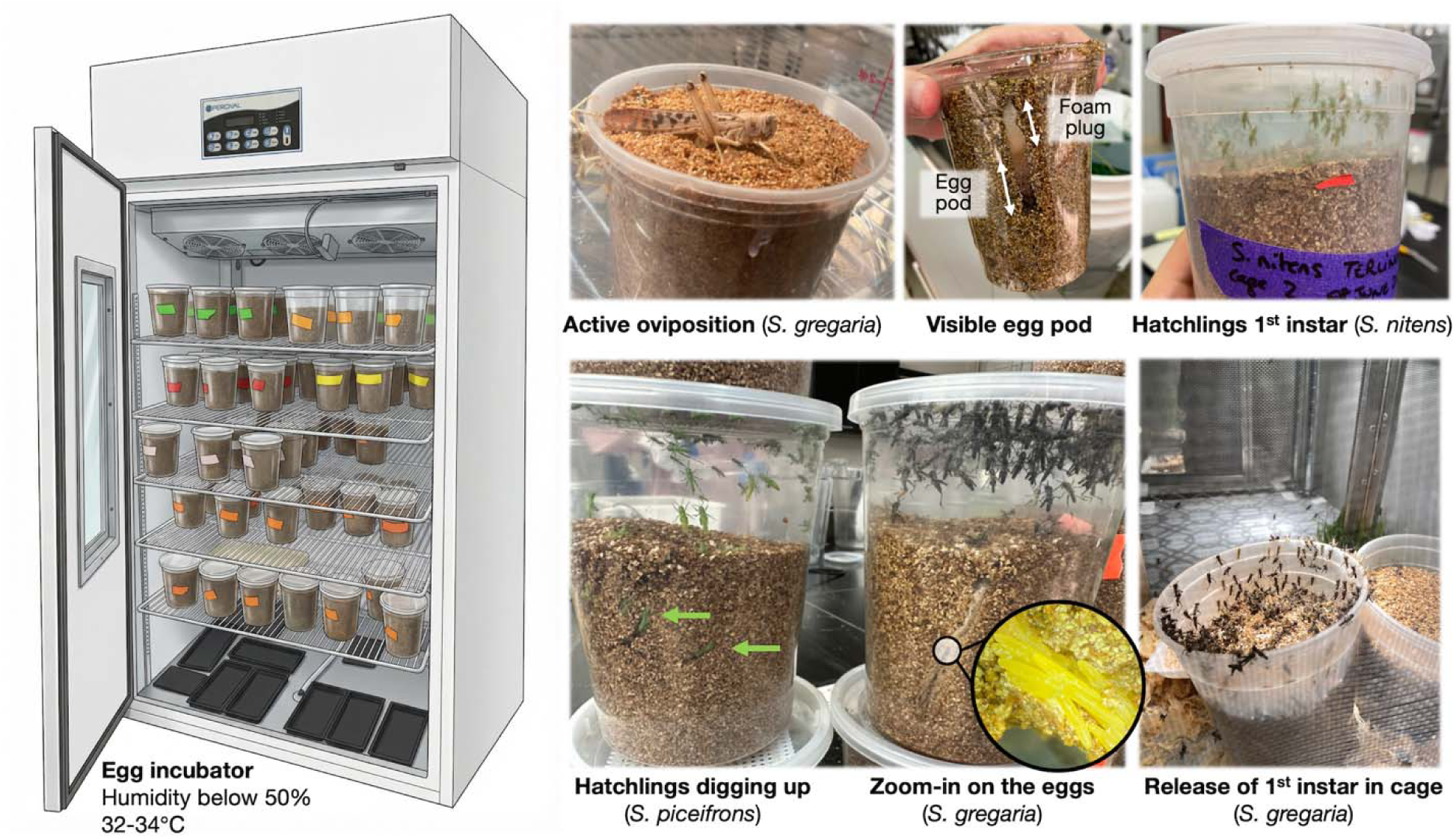
Process for egg pod inspection, incubation, and hatchling release across *Schistocerca* species. A single incubator can host multiple species, provided cups are clearly color-coded and labeled with species and oviposition date. Following active oviposition in the vermiculite, egg pods are identified by the presence of a foam plug and egg cluster, which may be visible at the surface or along the cup wall. Hatchlings emerge by crawling upward through the vermiculite.

**Figure 7:**
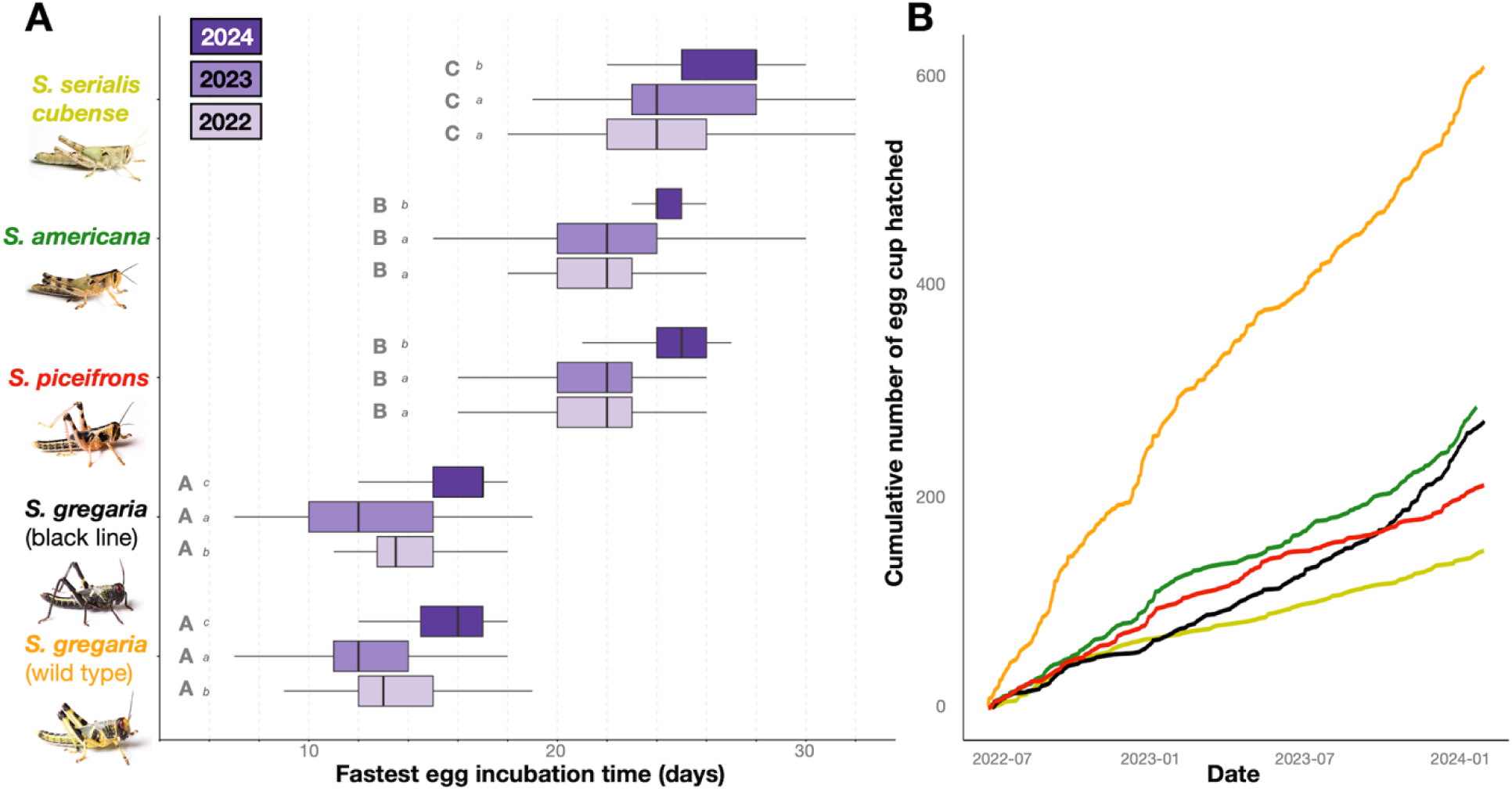
Incubation time and hatching dynamics for different *Schistocerca* species and laboratory lines maintained in long-term crowded high density. (A) Egg incubation time was measured from a total of 1,530 egg cups collected between 2022 and 2024 at the Texas A&M University Locust Quarantine Facility, including *S. gregaria* wild type (n = 609) and black line (n = 273), *S. piceifrons* (n = 212), *S. americana* (n = 286), and *S. serialis cubense* (n = 150). *S. gregaria* eggs typically hatched within ∼14 days, whereas the slowest species, *S. serialis cubense,* required ∼20–25 days. Type II ANOVA revealed significant effects of Species, Year, and Species × Year interaction (***, ***, ***). Lowercase letters indicate Tukey post hoc comparisons among years within species; uppercase letters indicate Tukey post hoc comparisons among species within year. (B) Cumulative number of hatched egg cups.

**CRITICAL:** Avoid mixing late and early instars at high density, as this may lead to cannibalism and higher mortality, especially during molting.

e. Repeat transfers with successive egg cups until the desired density or third-instar nymphs appear.

**NOTE:** Propagation can be rapid for locust species using the same egg cup collection time window, such as *S. gregaria* (see Fig. 7B).

**7. Track the species life cycle**

a. Use standardized high-quality life-cycle monitoring sheets, adapted from Texas A&M University stock populations, to identify instars (Fig. 8).
b. Identify molt stages based on exuviae and changes in body size and coloration.
c. Alternatively, the instar stage can be approximated from the number of eye stripes. A new vertical stripe is formed after each molt, ^24,25^ (Fig. 9).
**8. Stabilize colonies before experimental use.**

a. Rear colonies under standardized common-garden conditions for a minimum of two generations before initiating experiments.
b. Confirm that colonies reach a stable breeding state, defined by sustained egg pod production following sexual maturity in cages containing 50–100 adults (typically 2–3 weeks after adult molt).
c. Verify egg pod viability by confirming consistent hatching success (Fig. 7B), with approximately 20–60 hatchlings per egg pod (Fig. 6), depending on species.
d. Monitor development time, survival, and reproductive output across generations to confirm colony health and consistency.
e. Initiate experimental density manipulations and sampling only after stable colony performance is achieved.

**Figure 8:**
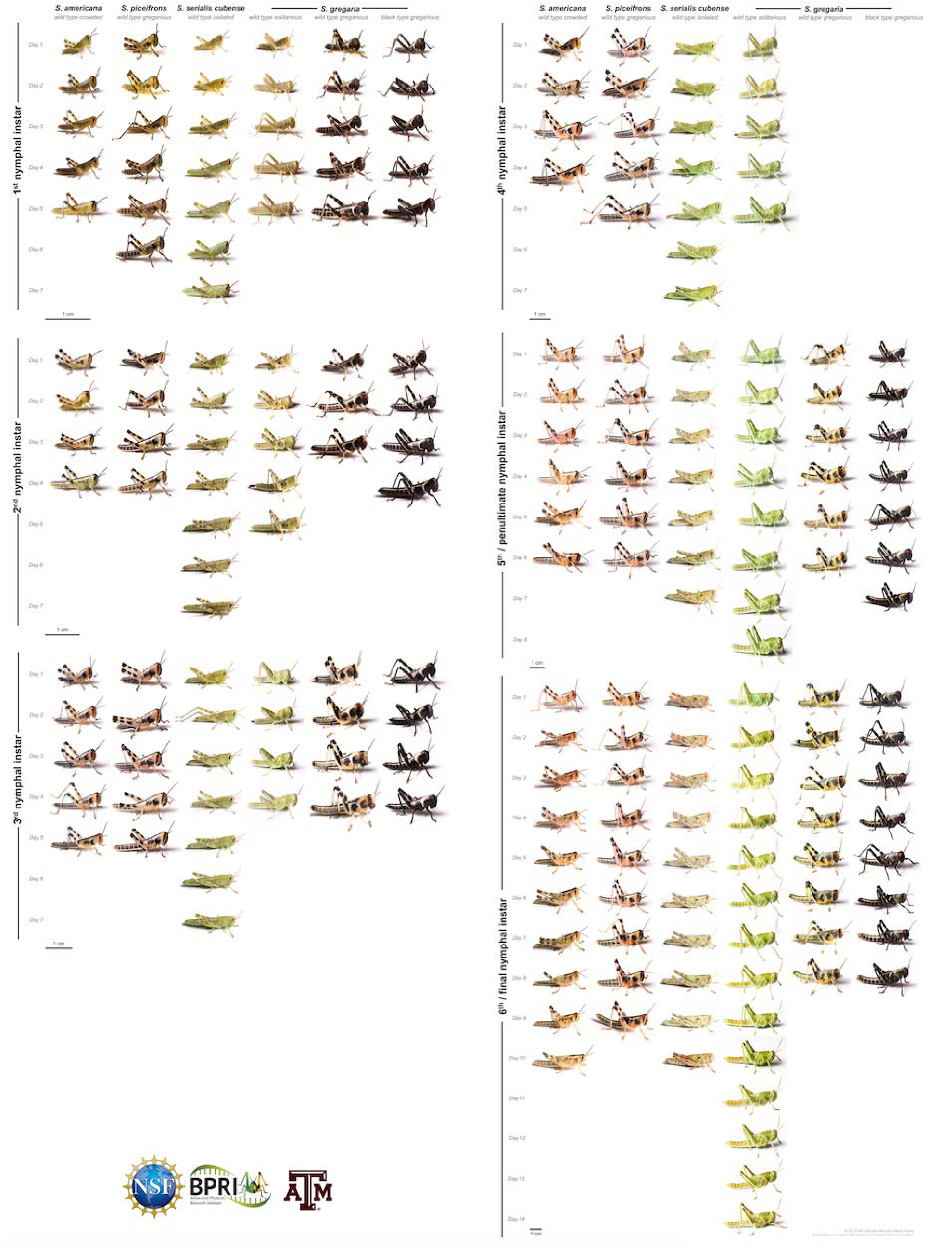
Nymphal developmental identification sheet generated from high-resolution daily imaging under controlled rearing conditions. This reference panel was built by rearing up to 100 nymphs in clear containers under common-garden conditions and feeding *ad libitum* with wheatgrass, lettuce, and wheat bran. Cages were cleaned and inspected daily for exuviae to accurately track molts. For each species and density condition (*S. gregaria* wild type and black line crowded, *S. gregaria* wild type isolated, *S. piceifrons* crowded, *S. americana* crowded, and *S. serialis cubense* isolated), lateral and scaled dorsal photographs were taken daily using a portable insect imaging whitebox studio setup using a Canon EOS Rebel T3 with an attached Canon 100 mm macro lens, and an external flash (Sunpak Auto 383 Super connected with a CowboyStudio 4 Channel Wireless Hot Shoe Flash Trigger & Receiver). Individuals were followed longitudinally when possible; in highly crowded cages, the fastest-growing nymph was selected to represent each instar. This dataset provides a standardized visual guide for instar identification across species and density regimes. Credits: Brandon Woo.

**Figure 9:**
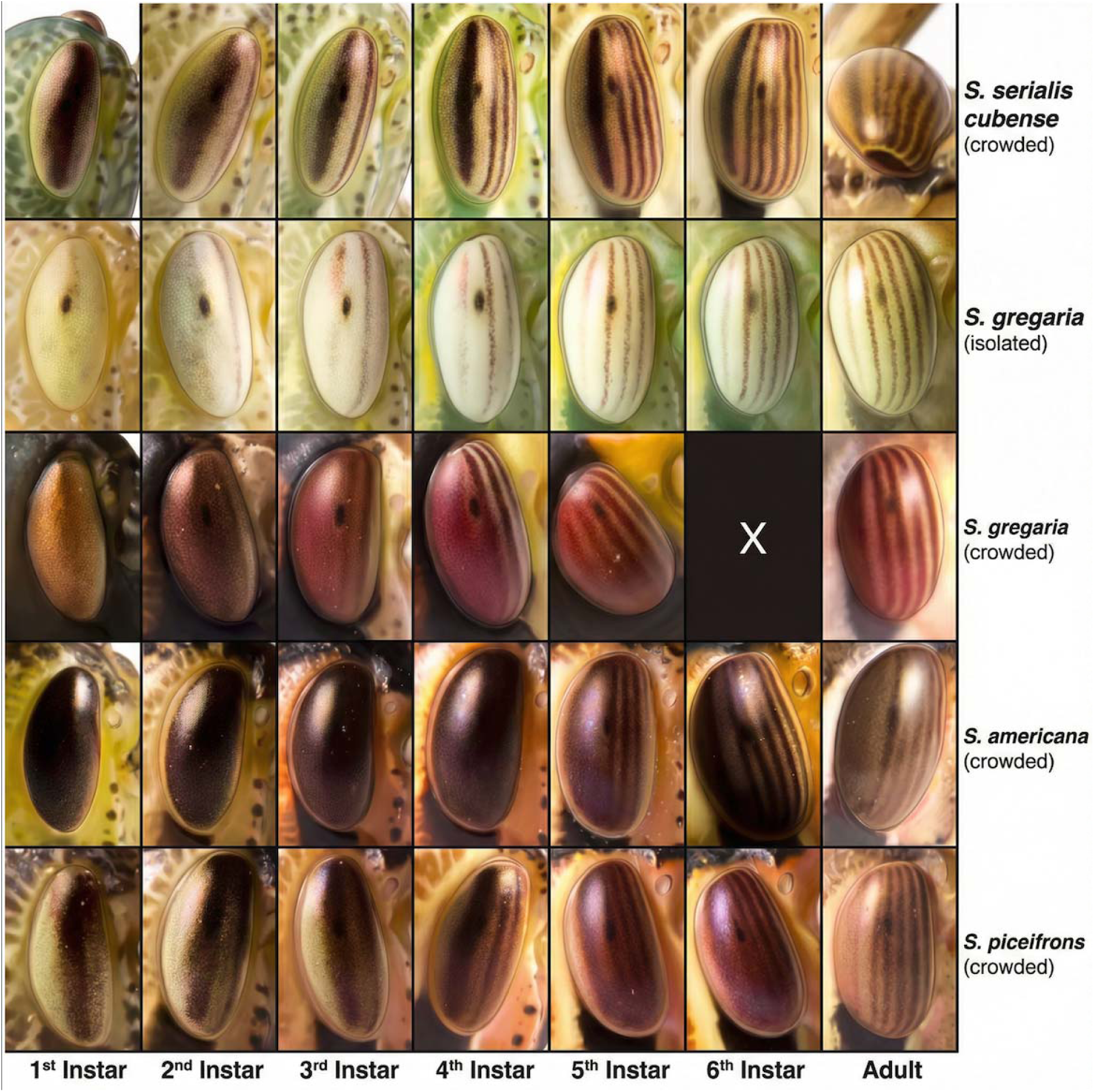
Eye stripe-interstripe patterns as an external marker of instar progression. The number of eye stripes and interstripes follows an approximately one-molt–one-stripe relationship and can be used to identify the instar stage. This method is reliable in *S. gregaria* and *S. serialis cubense* but is more difficult to apply in early instars of *S. piceifrons* and *S. americana*, where stripe contrast and definition are reduced. Eye pictures zoomed-in views of the specimens shown in Figure 7.

**NOTE:** Species with prolonged or facultative diapause (e.g., *S. nitens*, *S. piceifrons*) may require additional generations to stabilize.

### Part 2: Crowded and isolated rearing

**Timing:** 30 days (variable depending on species, instar targeted, and experimental design)

This step describes how to rear *Schistocerca* individuals under controlled, crowded, or isolated conditions to induce and maintain gregarious or solitarious phenotypes before behavioral assays, time-course sampling, or tissue dissection.

**NOTE:** For transcriptomic experiments in our laboratory, the reference stage is typically female last instar nymphs, 3 days post-molt, which is developmentally larger and identifiable across species and avoids inconsistencies caused by variable instar number (e.g., fifth instar skipping in *S. gregaria*).

**1. Initiate experimental cohorts**

a. Collect laboratory-laid egg pods from stabilized long-term crowded colonies for the species of interest, as described in Part 1.
b. Incubate egg pods under standard common-garden conditions until hatching.
c. Optionally use hatchlings from the same egg pod or from synchronized egg pods to generate both crowded and isolated cohorts, thereby minimizing genetic and maternal effects.
**2. Establish crowded rearing (gregarious condition)**

a. Select hatchlings or early instar nymphs of the same developmental stage (if originating from highly crowded cages).
b. Transfer individuals into a clean, pre-assembled, crowded cage (see Preparation of locust and grasshopper cages) to achieve high density.

i. Typical experimental densities range from 100 to 300 nymphs per cubic cage, depending on species and cage type.
**3. Establish isolated rearing (solitarious condition)**

a. Allow hatchlings from the same egg pod to emerge in a temporary container with fresh food for ∼24 h.
b. Prepare isolated cages in advance with food and ventilation components installed.
c. Transfer individuals one at a time into isolated cages.

i. Open the front door 1–2 cm above the gray tray surface.
ii. Use a Falcon tube or a similar container to guide each nymph into the cage, avoiding direct handling or crushing.
iii. House only one individual per cage throughout development.
iv. Close the front door tightly after successful introduction and label the cage with species and date.
d. Ensure complete visual, tactile, and olfactory isolation from conspecifics.

i. Verify that the airflow for the tubing system is above 9 LPM.
ii. Check that the activated charcoal filter is installed before use (Fig. 2B).
**4. Implement standardized feeding under crowded and isolated conditions**

a. Provide food *ad libitum* throughout the development.

i. Replace wilted or contaminated plant material immediately.
ii. Refill wheat bran daily in a clean Petri dish.
iii. Under high-density crowded conditions, feed up to twice daily to reduce food competition and cannibalism, particularly during molting.
iv. Present fresh plant material upright in a glass jar filled with water, using organic romaine lettuce, wheatgrass, and wheat bran as the primary sources of food.
v. Add sufficient plant material to prevent young nymphs from falling into standing water.
vi. Position food slightly offset from the central heat/light source to reduce fecal accumulation and localized moisture buildup.
**5. Clean cages and maintain hygiene**

a. Clean crowded cages frequently by scraping accumulated frass or adding a fresh layer of wood shavings as needed.
b. Clean isolated cages according to developmental stage.

i. For 1st–4th instars, remove frass every 3-4 days using a handheld vacuum fitted with flexible PVC tubing.
ii. For 5th instar nymphs and adults, remove frass daily or every 2 days.
**6. Monitor phenotype expression and health**

a. Maintain standard environmental conditions (30 °C, 12 h light:12 h dark).
b. Inspect cages daily for mortality, mold, or ventilation issues.
c. In isolated individuals, monitor reduced activity, increased wariness, and avoidance behaviors characteristic of solitarious phenotypes
d. If needed, record developmental timing and molting events to ensure accurate stage matching across treatments.

**NOTE:** Species like *S. americana* and *S. piceifrons* can still show high escape-avoidance activity, such as jumping, in the solitary phase.

### Part 3: Behavioral assays

**Timing:** 1h for arena setup, 15 mins for each assay

This experiment is performed using a rectangular Roessingh-style arena (57 x 31 x 11 cm) for late instars/adults or a smaller version (35 x 15 x 7.5cm) for first instars, to evaluate locomotor activity and attraction to conspecifics. Walls of the arena are made of mat Perspex, and the separations between the central compartment and the side chambers are made of clear Perspex with holes. The two side chambers are also made of clear Perspex, with one face perforated.

This setup uses a camera mounted above the arena to record behavioral activity. At Texas A&M University, it was achieved by constructing a steel frame lined externally with white shower-curtain material, secured to the frame with glued magnets. The arena is positioned directly beneath the camera, ensuring the central compartment is fully visible. Video recordings are subsequently used to track the focal individual’s behavior using behavioral tracking software; Ethovision XT versions 12 and 17 were used at Texas A&M University.

1. Fill one of the side chambers with 50 final instar male and female nymphs. Keep the other one empty.

**NOTE**: Before starting the experiment, one should ensure there are enough non-focal individuals available as stimuli. A couple of young adults can be used to reach 50.

2. Place a focal final instar nymph marked 72 hours prior (see Part 4 Step 4) in a blackened syringe for approximately 2 minutes to reduce handling stress.

1. **NOTE**: Prior to the experiment, a 50mL Falcon tube needs to be lined with black tape, and a syringe plunger needs to be added so the grasshopper can be ejected.
3. Insert the nymph through the bottom hole in the middle of the arena using the syringe, and keep the plunger inserted so the hole is covered.
4. Close the curtain, start the recording in Ethovision XT, and let it run for 12 minutes.
5. After 12 minutes, end the recording trial and prepare for the next one:

1. Remove the plexiglass cover
2. Retrieve the focal nymph by hand and discard it as needed, depending on the experiment (freeze it, return it to its original cage, set it aside, etc.).
3. Clean the arena with 70% Ethanol or with a disinfectant wipe.
4. Start again with the next focal nymph.
6. Once all trials are run, place the 50 stimulus conspecifics back in their original cage, and clean the chamber and the entire arena with 70% Ethanol or a disinfectant wipe.

**NOTE:** If nymphs require dsRNA injection, the protocol remains unchanged, except for trial timing. Once newly molted final instar nymphs are marked with paint pens, they are food-deprived for 24h before injection. Once injected with the dsRNAi of choice, they are returned to their cage with food ad libitum until the behavioral trial, 72 hours post-injection.

### Part 4: Time-course sampling design

**Timing:** 3-7 days (variable depending on sampling scheme)

This step describes how to perform controlled time-course experiments to track molecular responses before, during, and after density-dependent phase transitions in *Schistocerca*. The protocol emphasizes precise timing, individual marking and identification, and synchronized handling across treatments to accurately capture rapid and delayed transcriptional responses associated with phase change. For transcriptomic studies, include at least 5 biological replicates per condition and time point to capture inter-individual variability and disentangle shared gene expression dynamics associated with density-induced phase transitions.

**NOTE:** For transcriptomic experiments in our laboratory, the reference stage is typically female last instar nymphs, 3 days post-molt, which is developmentally larger and identifiable across species and avoids inconsistencies caused by variable instar number (e.g., fifth instar skipping in *S. gregaria*).

**1. Define the density transition to be tested and prepare dedicated housing.**

**a.** Prepare material for gregarization: the transfer of individuals from isolated to crowded conditions.

i. A stimulus clean cubic cage with 200-500 nymphs of the same instar should be available and dedicated to the experiment.
**b.** Prepare material for solitarization: the transfer of individuals from crowded to isolated conditions.

i. Prepare empty, clean, fully assembled isolated cages in advance (see Preparation of locust and grasshopper cages).
ii. Verify airflow, charcoal filter placement, and door sealing before starting the isolation process.
**2. Define experimental time points and a standardized start window.**

**a.** Include a t0 control sampled directly from the long-term baseline condition immediately before transfer.
**b.** Sample short-term responses at 30 min to 4 h after transfer. Use these time points to capture rapid neural and behavioral-response-associated transcriptional changes.
**c.** Sample intermediate responses at 4 to 72 h after transfer. Use these time points to capture sustained regulatory shifts as individuals acclimate to the new density context.
**d.** Sample long-term phase responses at >72 h after transfer. Use these time points to capture stabilized phase-associated expression profiles under the new density condition.
**e.** Define a fixed daily start window for density changes (e.g., perform all introductions between 09:00 and 12:00) to reduce circadian and handling variability.
**3. Select the target developmental stage and sex for the time-course experiment.** **NOTE:** Here, we target female last-instar nymphs 3 days post-molt (Fig. 8).
  **a.** Confirm last instar status using species-specific instar cues (e.g., body size, wing pad development), with reference to the developmental cheat sheet (Fig. X).
  **b.** For crowded colonies, visually confirm that candidate cages contain ≥ 200 last-instar nymphs before cohort selection.
  **c.** If the experimental crowded cohort contains fewer than 100 individuals, supplement with younger instars from synchronized stock cages as needed.
  **d.** For isolated individuals, sex individuals at the target stage to ensure sufficient females for all planned time points and biological replicates (Fig. 5).
**4. Standardize molt tracking before initiating density transitions using a cut-off system and color coding.**

**a.** Define a 12-h or 24-h cut-off window for molt assignment based on manpower and cohort size.
**b.** Begin tracking when the first last-instar molts are observed in the experimental cohort cage.
**c.** Remove non-tracked last-instar individuals from the cohort cage to reduce staging ambiguity. Alternatively, keep the individuals in the cage but by marking with a specific color.
**d.** Assign a unique Posca paint pen color (or combination) to each cut-off window or tracking day and record the color–date/time mapping.
**5. Track, sex, and sort newly molted crowded candidates during each cut-off window.**

**a.** Inspect the cohort cage every 12 h (typically during feeding) for 30–40 min under bright light.
**b.** Check egg carton folds and other hiding sites where newly molted nymphs aggregate.
**c.** Transfer newly molted last-instar nymphs gently into a temporary sorting container by separating sexes.
**d.** Mark individuals with Posca Paint Pens and allow the paint to dry (Fig. 10).

i. Use a clear container with perching to prevent individuals from walking over one another during paint drying (5-10 min), or
ii. If needed for highly active insects, slice a sponge foam tray (used for packing or jewelry box lining) to create a tray to briefly immobilize individuals for marking and drying.
iii. Mark non-experimental individuals (males and damaged/abnormal females) with an exclusion color (e.g., white).
iv. Mark suitable experimental females with the assigned cut-off color and record the date/time window and number marked.
**e.** Return marked individuals to their designated baseline condition cage and provide food *ad libitum* with egg carton perches (Fig. 10).
**6. Repeat Step 5 until sufficient females are staged within the same cut-off window to run parallel transitions and sampling.**
**7. Track and mark isolated individuals by direct observation.**

**a.** Inspect each isolated cage every 12 h for exuviae and record the molt date/time window.
**b.** Mark the nymph with the assigned cut-off color and return it immediately to the same isolated cage.
**c.** Provide food *ad libitum* throughout staging.
**8. Initiate density transitions using age-matched females and begin the time course.**

**a.** Start the density change when staged females reach the target age (e.g., **3 days post-molt**) and within the standardized introduction window defined in Step 2e.
**b.** For **gregarization**, transfer one isolated marked nymph into the stimulus-crowded cage for the desired exposure duration (Fig. 10).

i. Transport the nymph in a small deli cup and introduce it through the stockinette sleeve.
ii. Provide additional food if needed during high-density exposure.
iii. Optionally monitor behavior toward conspecifics and food at defined intervals without disturbing the cage.
iv. At the end of the time point, collect the marked nymph with gloved hands and place it into a clean deli container with small ventilation holes.
v. Immediately proceed to dissection or snap-freezing (as appropriate for downstream assays).
**c.** For **solitarization**, transfer one marked crowded nymph into a prepared isolated cage for the desired exposure duration.

i. Transport the nymph in a small deli cup and introduce it through the front door, then close the door tightly.
ii. Provide additional food if needed.
iii. (Optional) Monitor food and conspecific-related behavior at defined intervals without excessive disturbance.
iv. At the end of the time point, collect the nymph with gloved hands and place it into a clean deli container with small ventilation holes.
v. Immediately proceed to dissection or snap-freezing (as appropriate for downstream assays).

**Figure 10:**
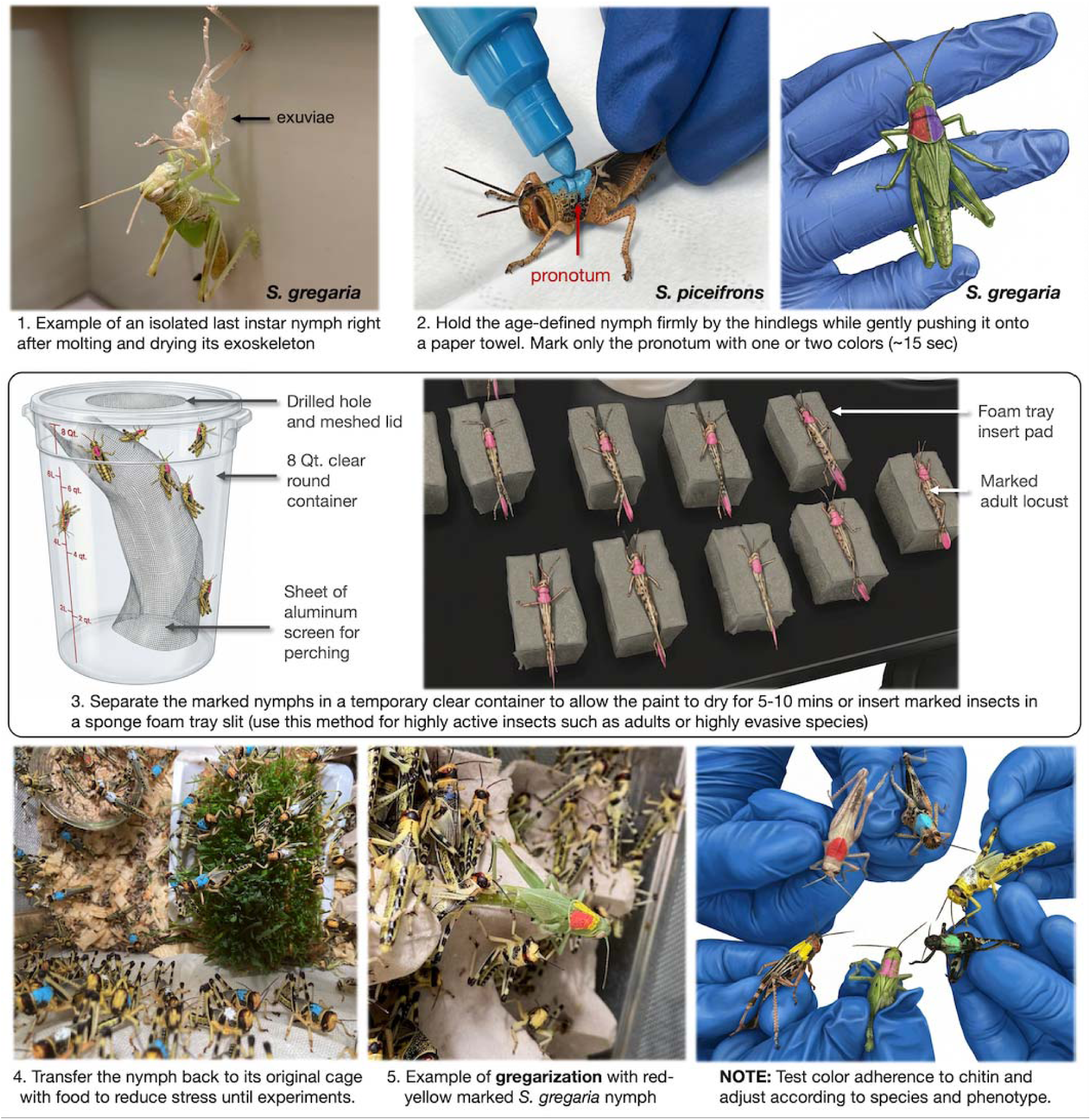
Schematic and photographic overview of pronotum marking using non-toxic water paint pens to identify age-defined nymphs. Individuals are briefly restrained, marked on the pronotum, allowed to dry in a ventilated temporary container or on a foam pad, and then returned to their original cage. When applied correctly, color can adhere to the nymph for several days and is eliminated during the molting process.

**NOTE:** Molt tracking is straightforward in isolated cages because each individual can be scored directly. In crowded cages, staging is less reliable without temporary standardization, because exuviae are quickly trampled, eaten, or mixed among many individuals, and newly molted individuals can be difficult to re-identify without controlled observation.

### Part 5A: Dissections of nymph nervous, chemosensory, and digestive tissues

**Timing**: 5–30 min per individual (depending on tissues collected)

This step describes the rapid, sequential dissection of whole locust or grasshopper bodies to isolate head-, gut-, thoracic-, nervous-, and chemosensory-associated tissues from a single individual while minimizing contamination, RNA degradation, and handling-induced transcriptional artifacts. The procedure is optimized for last-instar female nymphs but can be adapted to late instars and males (4th–6th instar).

We recommend live tissue processing for RNA extraction. Although storage in ethanol or RNAlater is more convenient, our extensive testing revealed that even overnight storage substantially complicates tissue separation after thawing.

**Glove requirement:** All steps require clean nitrile gloves. If glove contamination is suspected (e.g., regurgitation or fresh defecation), spray and rub the gloves with 70% ethanol, or change them immediately.

**NOTE:** Use multiple sets of forceps and scissors to minimize cross-contamination between tissues. Hot-bead sterilization between steps is possible, but allow tools to cool fully before reuse to avoid RNA degradation. Avoid idle time between steps.

Separate the head, thorax, and abdomen.

**Timing**: 5 min

**1. Retrieve, inspect, and immobilize the specimen.**

a. Gently remove the individual from the transfer container, avoiding deformation of the abdomen or thorax.

i. Confirm sex by examining the subgenital plate and valvulae (Fig. 5).
ii. Record any visible damage or deformities resulting from experimental treatment.
iii. *(Optional)* Take a rapid photograph of the individual to document body coloration or markings.
b. Place the specimen ventral side up on clean aluminum foil, with the legs oriented toward you.

i. Hold the pronotum steady by hand and immobilize the specimen (Fig. 11A).
ii. Using large spring scissors, cut all legs at the coxae (bases attached to the thorax).
iii. (Optional) Collect hemolymph from the leg wounds within ∼10 s using capillary tubes or a 50 µL pipette with a filtered tip; lift the specimen during collection to prevent hemolymph from contacting the aluminum foil (Fig. 11B).
c. Remove external fluids and position the specimen.

i. Lift the legless specimen by the pronotum using large forceps
ii. Gently blot the ventral surface on a Kim wipe to remove regurgitation reflex and excess hemolymph.
iii. Return the specimen ventral side up onto a clean area of aluminum foil.
**2. Detach and stabilize the head.**

a. Hold the pronotum laterally with large forceps or with a clean glove, avoiding compression of internal tissues.
b. Using large dissection scissors in the open position, pull the head gently away from the thorax to expose the cervix soft junction, ensuring maxillary and labial palps are not obstructing the cut (Fig. 11C).
c. Sever the head in one swift, decisive motion to prevent foregut leakage and external contamination.
d. Immediately transfer the head to a chilled dissection plate and fully submerge it in ice-cold locust saline.

**Figure 11:**
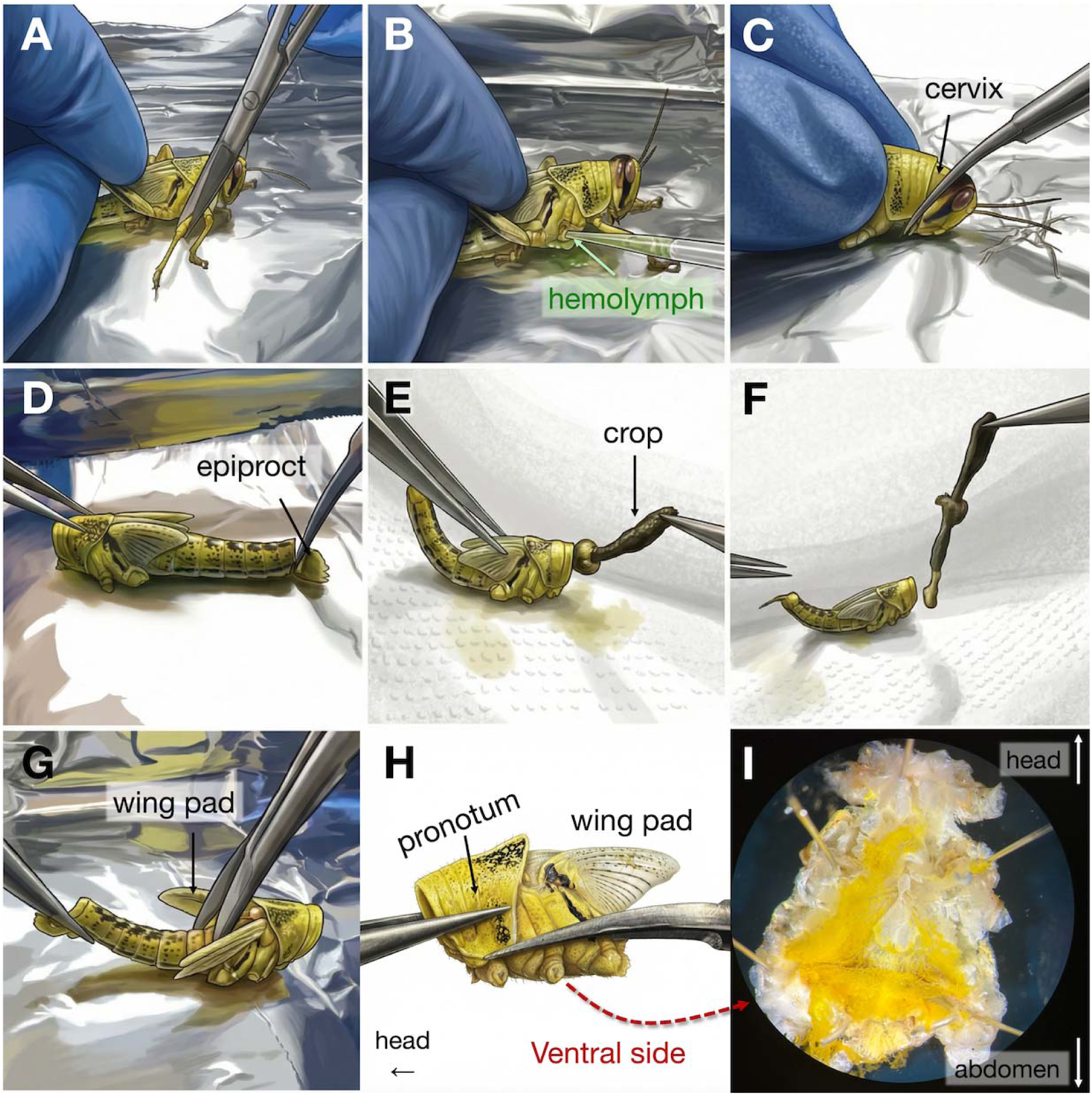
Key steps for rapid separation of head, gut, thorax, and abdomen during whole-body dissection. (A–C) Immobilization, hemolymph collection, and head detachment at the cervical membrane. (D–F) Abdominal incision and extraction of the intact gut by grasping the crop. (G–H) Separation of the thorax and abdomen at the wing pad junction and isolation of the ventral thorax. (I) Ventral view of the pinned thoracic cavity showing internal organization without cleaning. Labeled anatomical landmarks highlight reproducible reference points used to minimize tissue damage and cross-contamination.

**CRITICAL**: If necessary, pin the head above the clypeus to keep it fully submerged while avoiding damage to the central nervous system. Do not leave the head unprocessed for >10 min to prevent RNA degradation and stress-induced transcriptional changes.

For the central nervous system, palps, or antenna dissections only, proceed directly to the corresponding downstream steps (see Step 6).

**3. Release and extract the alimentary canal.**

a. Wipe the tips of the dissection scissors with 70% ethanol and a Kimwipe.
b. While holding the pronotum gently with large forceps, cut vertically through the last abdominal tergite above the epiproct to release the rectum (Fig. 11D).
c. Immediately place the large scissors face down into a hot ddH₂O bead beaker with a silicone bottom to prevent cross-contamination.
d. Using a new, clean pair of forceps, firmly grasp the exposed crop-foregut at the head-side opening and pull the gut upward in one continuous motion, while holding the pronotum with the previously used forceps (Fig. 11EF).
e. Transfer the intact alimentary canal immediately to a separate chilled dissection plate containing ice-cold locust saline (**see Step 11**), or directly into a labeled tube for storage.

**CRITICAL:** Do not allow the alimentary canal to come into contact with the aluminum foil. A poor grip on the crop may tear the membrane and release bacterial contents, as the junctions between the foregut, midgut, and hindgut are soft and fragile.

f. Place the forceps and scissors used for gut extraction face down into the hot ddH₂O bead beaker with a silicone bottom before proceeding to subsequent dissection steps.

**4. Separate thorax and abdomen.**

a. While holding the pronotum with the previously used large forceps, take a new large dissection scissor in the closed position and slide the tips between the wing pads to their base on the first abdominal tergite.
b. Open the scissors gently to spread the wings and cut vertically through the thorax–abdomen junction in one swift, strong motion (Fig. 11G).
c. Place the detached abdomen into a labeled tube designated for body leftovers, if required.
d. Clean the forceps used to hold the pronotum with 70% ethanol and a Kimwipe.
e. Using a new, clean pair of forceps, grasp the inner lateral side of the pronotum at the head–thorax cut site and lift the thorax from the aluminum foil.
f. Cut laterally above the leg coxae and below the wing pads to separate the dorsal and ventral thoracic sections (Fig. 11H).
g. Tear dorsal and ventral sections apart and immediately submerge and pin the ventral thorax in an ice-cold saline dissection plate (Fig. 11I).
h. Place the dorsal thoracic cuticle into the labeled tube for body leftovers, which is kept on ice.

### Dissection of thoracic tissues

**Timing**: 5 min

**5. Dissection of the metathoracic ganglia.**

a. Take the dissection plate containing the ventral thoracic section, submerged in ice-cold locust saline solution.
b. Orient the thorax so that the head–thorax cut edge is facing upward and the thorax–abdomen cut edge is facing downward to differentiate the ganglia (Fig. 12AB) properly.
c. Pin the four extremities of the thoracic cuticle so the tissue is fully stretched and flat.
d. Place the forceps used for thorax handling into the hot ddH₂O beaker with a silicone bottom for cleaning.
e. (Optional) Using a new pair of fine forceps (Dumont #5), collect 5–20 mg of fat body lining the inner thoracic cavity (Fig. 12C).

i. i. Identify fat body as a yellow, amorphous, slightly granular tissue attached to the cuticle and muscle.
ii. Transfer the fat body into a new, labeled tube and snap-freeze.
iii. Place the forceps used for fat body collection into the hot ddH₂O beaker for sterilization.
f. To expose the meso- and metathoracic ganglia, remove air sacs, remaining fat body (Fig. 12C), thoracic muscle (Fig. 12D), air sacs and trachea (Fig. 12E) using fine forceps (#DF5A), discarding tissue onto a Kim wipe.

i. If visibility is reduced, gently flush the cavity with ice-cold locust saline to clear debris and improve visualization.
ii. Remove excess of locust saline in a waste cup if needed.
g. Once the ganglia are exposed, use fine dissection scissors to cut the nerve cords between the mesothoracic and metathoracic ganglia (Fig. 12F).
h. Inspect the isolated metathoracic ganglion using fine forceps (#DF5A) and remove any remaining tracheae or air sacs (Fig. 12G).
i. Briefly rinse the metathoracic ganglion by gently swooshing it in the locust saline away from the thorax.
j. Transfer the ganglion into a labeled screw cap tube and snap-freeze before storing at -80℃ for RNA processing.

**Figure 12:**
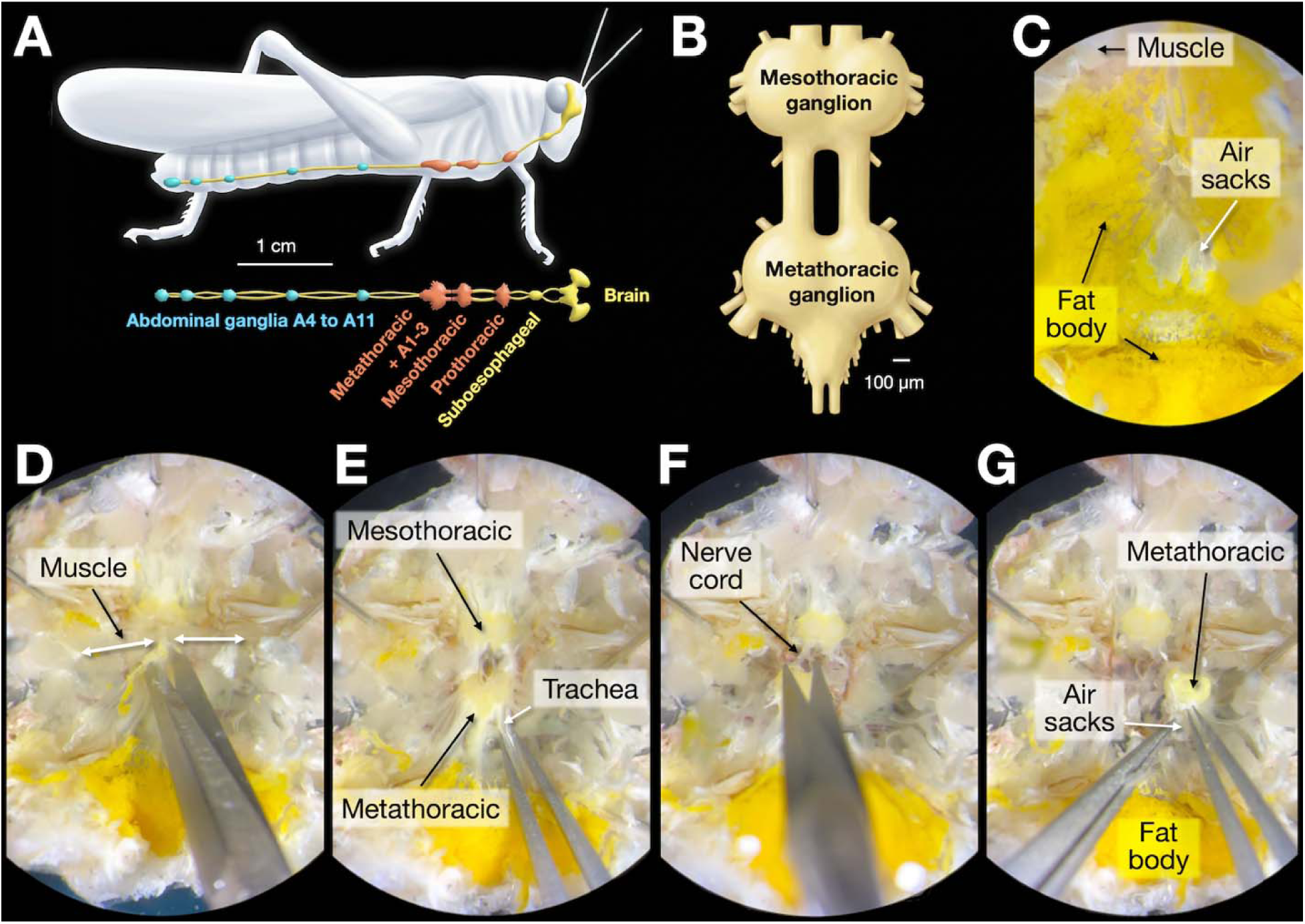
Anatomy and key steps for the isolation of the metathoracic ganglia during ventral thorax dissection. (A) Schematic overview of the locust central nervous system, highlighting the brain, suboesophageal ganglion, thoracic ganglia, and abdominal chain (adapted from ^26^). (B) Expected morphology of the metathoracic ganglia in comparison with the mesothoracic ones. (C–G) Sequential views during ventral thorax dissection showing removal o muscle, fat body, air sacs, and tracheae to expose the metathoracic ganglia and connecting nerve cord.

### Dissection of antennas, maxillary palps, and brain

**Timing**: 12-15 min

**6. Isolate chemosensory-associated structures and the brain.**

a. Using a large forceps (Pelco) and a large spring scissor, stabilize the head submerged in ice-cold locust saline and prepare the cut above the mouthparts.

i. Use the scissors to gently push the floating head downward.
ii. Hold the preocular ridge along the vertex with forceps while keeping the head submerged.
iii. If the grip is unstable, temporarily remove excess saline to improve handling.
b. With scissors in the open position, place the blades just at the clypeus–mandible junction (Fig. 13AB).
c. In one swift, strong, and decisive motion, cut the mouthparts from the rest of the head (Fig. 13B).

i. Expect an audible cracking sound due to thick chitin.
ii. If the mouthparts fragment, carefully collect all pieces, including palps. **CRITICAL:** Do not cut too much above the line, as it may damage the central nervous tissue.
d. Using forceps, transfer the entire mouthpart complex (clypeus, mandibles, labrum, maxillae, labium) into a labeled screw cap tube and snap-freeze before storing at -80℃ for RNA processing.
e. *(Optional)* Perform further fine dissection to isolate maxillary or labial palps, retaining only the terminal 2–3 segments if required.
**7. Stabilization of the head for brain exposure.**

a. Pin the head securely in the dissection plate filled with ice-cold saline (Fig. 13C):

i. Insert pin #1 near the frontal costa along the vertex.
ii. Insert pin #2 behind the vertex above the compound eye (avoid inward angling).
iii. Insert pin #3 below the left compound eye on the gena plate.
iv. Insert pin #4 below the right compound eye on the gena plate.
b. Place the forceps and scissors, tip-down, in warm water for cleaning and temporary storage.
**8. Antennal dissection**

a. Rotate the dissection plate so the antennae face directly.
b. Using fine forceps that have been wiped with 70% ethanol, gently pull one antenna to expose its base.
c. Using a new fine pair of scissors, cut the antenna at its base (Fig. 13D).
d. Transfer each antenna into a labeled tube using forceps into a screw cap tube and snap-freeze before storing at -80℃ for RNA processing.
e. Wipe the tips of forceps and scissors with 70% ethanol before proceeding.
**9. Exposure of the brain in frontal view**

a. Using fine scissors, make shallow bilateral incisions below each compound eye along the gena junction (Fig. 13E). Do not cut deeply, ideally only the clear chitin.
b. Using a razor blade holder, gently incise the transparent facial chitin surrounding the compound eyes.

i. Join the bilateral incisions posteriorly by cutting behind pin #2 (Fig. 13F).
ii. Remove pin #1.
c. While holding pin #2 with fine forceps, use new fine forceps to grasp both sides of the frontal ridge near the antennal bases (Fig. 13G).
d. Slowly lift and remove the central facial cuticle (Fig. 13H).

i. Depending on the grip, this may remove only chitin or chitin plus attached muscle.
e. Using fine forceps #DF5C, expose the butterfly-shaped brain (Fig. 13I).

i. Carefully remove remaining muscle, air sacs, fat body, tracheae, and connective tissue, placing debris onto a Kimwipe.
ii. Flush the dissection site as needed with locust saline.
iii. Pour the liquid excess into the waste container and refill with clean locust saline.

**Figure 13:**
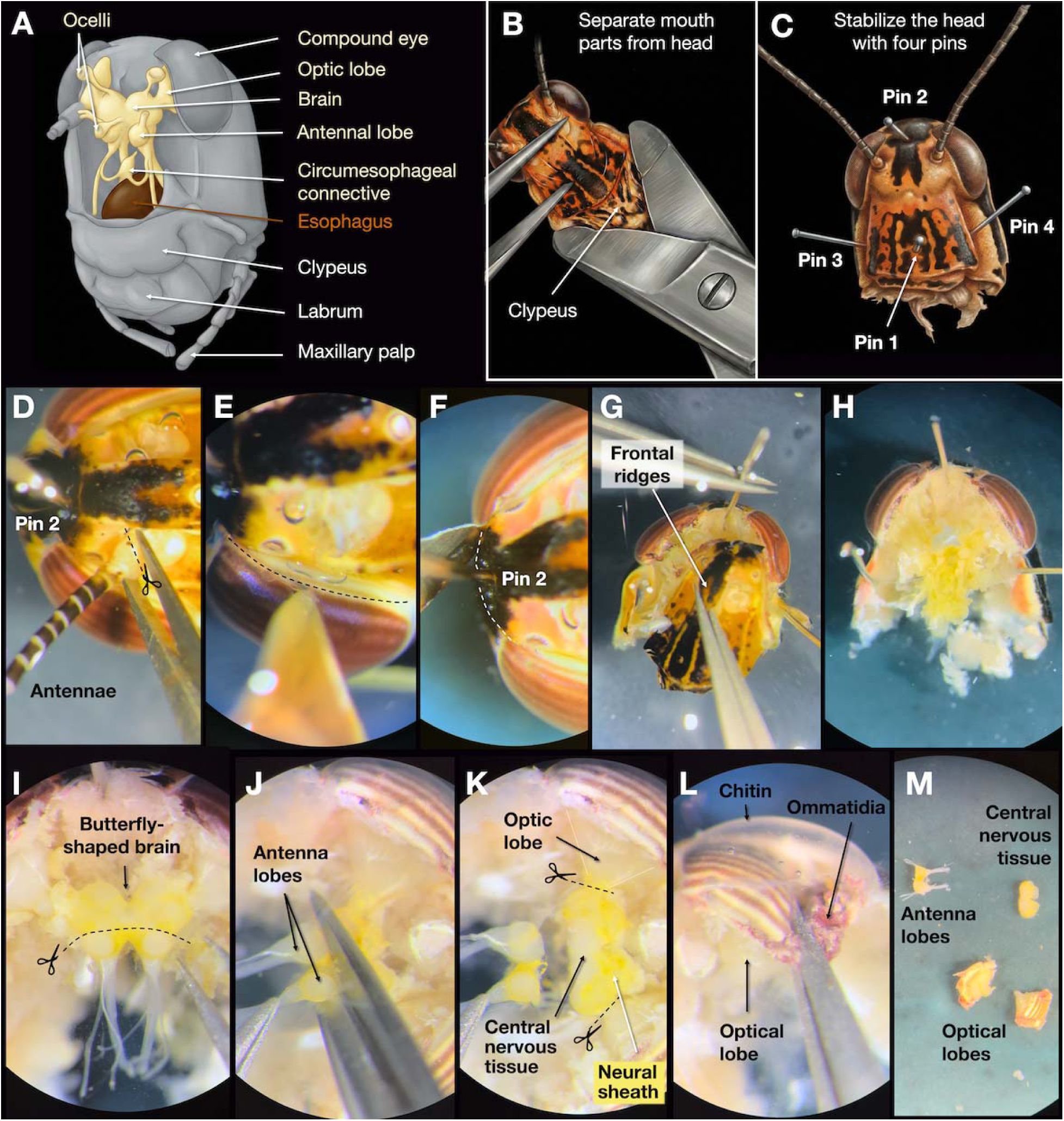
Anatomy and stepwise isolation of head-associated neural and sensory tissues. (A) Schematic overview of head anatomy highlighting the brain, antennal lobes, optic lobes, and mouthparts (adapted from ^27^) (B–C). Separation of mouthparts from the head and stabilization of the head using four headless pins. (D–H) Sequential incisions and removal of antenna, frontal cuticle, and ridges to expose underlying neural tissues. (I–K) Progressive exposure of the butterfly-shaped brain with antennal lobes and optic lobes following removal of overlying tissues and neural sheath. (L) Detachment of compound eyes and optic lobes from the surrounding chitin by scraping or grasping ommatidia. (M) Representative isolated head-derived tissues (central nervous tissue, antennal lobes, and optic lobes) are ready for downstream molecular analyses.

**CRITICAL:** Do not pull too strongly on the optic lobes junction as they can detach easily.

**10. Separate antennal lobes, optic lobes with compound eyes, and the central nervous system**

a. Using fine Dumont tweezers, gently remove the thin protective layer covering the brain.

i. This layer consists of fat body and neural sheath material and must be carefully peeled away.
ii. ii. Depending on species, this layer can be sticky and difficult to remove; *S. gregaria* is typically easier to process than *S. piceifrons*.
iii. iii. Avoid pulling directly on neural tissue while removing the sheath.
b. Using fine dissection scissors, gently grasp the nerve cords extending below the antennal lobes to slightly tension the tissue (Fig. 13J).

i. Cut just above both antennal lobes in a single, clean motion.
ii. Immediately transfer the paired antennal lobes into a screw cap tube and snap-freeze before storing at -80℃ for RNA processing.
c. Separate the remaining central brain (Fig. 13K).

i. Using fine scissors, sever the junction between the central brain and the optic lobes.
ii. Avoid compressing the optic lobes during cutting.
iii. Inspect the remaining central brain tissue and remove any residual protective sheath by gently peeling it away “like a sock” using fine tweezers.
iv. Transfer the cleaned central nervous tissue into a labeled screw cap tube and snap-freeze before storing at -80℃ for RNA processing.
d. Dissect optic lobes with compound eyes. Processing depends on the developmental stage after experimental treatment:

i. ≥6 days post-molt: the compound eye may detach from the chitinous eye capsule by gently pulling on the optic lobe.
ii. 3–6 days post-molt or if resistance is encountered: gently scrape away the outer ommatidial tissue from the chitin using fine tweezers to facilitate detachment.
iii. Holding the ommatidia, pull steadily to detach the optic lobe together with the compound eye (Fig. 13LM).
iv. Place the optic lobe + compound eye into a labeled tube and snap-freeze.
v. Repeat for the contralateral side and pool both tissues in the same screw cap tube and snap-freeze before storing at -80℃ for RNA processing.

### Dissection of the sectioned alimentary canal parts

**Timing**: 5 min

**11. Isolation of the midgut and caecum**

a. Pin the alimentary canal placed on a separate dissection plate at the start of the foregut and another at the end of the hindgut (following step 3 of Part 5A).
b. Perform a cut at the intersection of the proventriculus and the gastric caecum.
c. Lightly orient the gastric caecum towards the foregut and cut it from the midgut.
d. Brush back Malpighian tubules to orient them towards the hindgut.
e. Make a cut at the intersection of the ventriculus and ileum, fully removing the midgut.
f. Carefully remove any attached Malpighian tubules and trachea from the midgut.
g. Move the midgut to one well of a 24-cell well plate filled with cold sterile locust saline solution.
h. Make a midsagittal cut, then empty and wash the midgut into the well using tweezers.
i. Separate and place tissues in a labeled screw cap tube and snap-freeze in liquid nitrogen before storing at -80℃ for RNA processing.
**12. Hindgut**

a. Clean and wipe tweezers with ethanol and sterilize in hot microbeads for 10 s; or use a new pair of sterile tweezers.
b. Remove the pin from the hindgut and move it into a well plate filled with locust saline solution
c. Make a midsagittal cut, emptying and washing the hindgut into the well
d. Separate and place tissues in a labeled screw cap tube and snap-freeze in liquid nitrogen before storing at -80℃ for RNA processing.

### Part 5B: Dissections of adult female accessory gland and oviduct

**Timing:** 10–15 min per individual

This step describes the dissection of adult female reproductive tissues, with a focus on accessory glands and oviducts, from a single individual while preserving tissue integrity and minimizing contamination. The protocol is optimized for sexually mature adult females but can be adapted to late pre-oviposition stages.

**Glove requirement:** All steps require clean nitrile gloves. Change or ethanol-clean gloves if contamination is suspected.

**NOTE:** Prepare all pins and tools in advance to minimize handling time once the abdomen is opened.

**1. Prepare the dissection plate and immobilize the specimen.**

**a.** Insert ∼5 headless pins vertically into the silicone base of the dissection plate using large forceps.
**b.** Sterilize the forceps after use and return them to the tool tray.
**c.** Place the specimen on aluminum foil, hold it steady by the pronotum, and remove all legs and wings at the joints using large scissors (Fig. 14ABC).
2. **Position and pin the specimen** (Fig. 14D).

**a.** Transfer the specimen to the dissection plate.
**b.** Stabilize the thorax by pinning through the open leg joints.
**c.** Using fine tweezers, gently grasp the ovipositor and slightly extend the abdomen.
**d.** Insert a pin above the ovipositor plate to maintain tension.
**3. Open the abdominal cuticle.**

**a.** Using small scissors, cut longitudinally along one side of the abdomen from the thorax–abdomen junction to the ovipositor, following the spiracle line (Fig. 14D).
**b.** Repeat the cut on the opposite side.
**c.** Cut horizontally across the abdomen just posterior to the thorax to free the anterior edge of the cuticle.
**d.** Slowly lift the cuticle toward you, creating a hinged “flap.”

**Figure 14:**
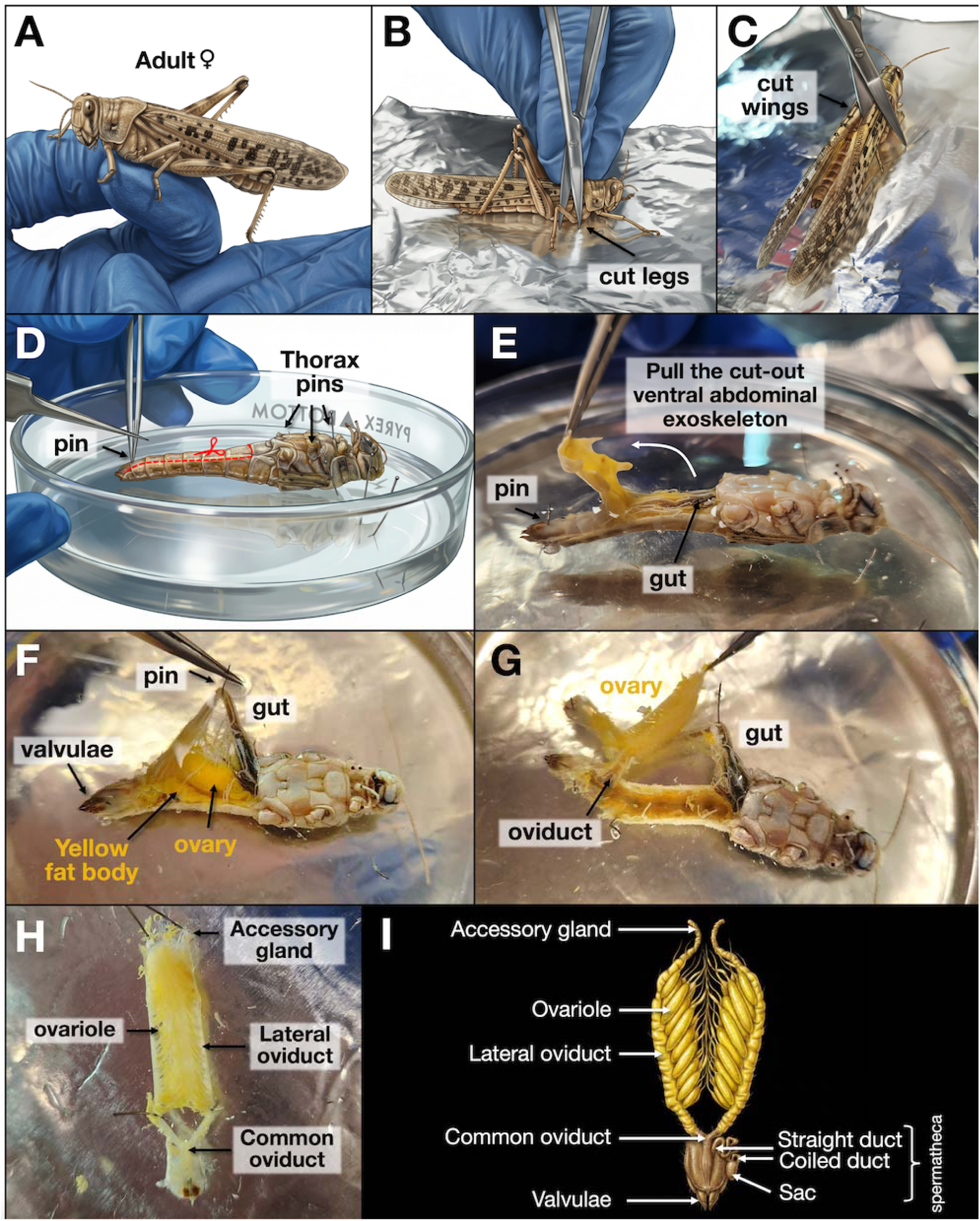
Anatomy and stepwise isolation of the female reproductive system of *Schistocerca*. (A) Handling of an adult female *S. gregaria*, held in place by the middle leg. (B-C) Separation of legs and wings to perform ventral dissection on an immobilized individual. (D-Sequential pinning, incisions, and removal of the abdominal ventral side exoskeleton to expose the alimentary canal, ovary, and fat body. (F-G) Overview of the internal structures before and after removing fat body, muscle, and connective tissues without damaging the gut and reproductive systems. Accessory glands can already be collected during step G without isolating the complete reproductive system. (H) Extraction out of the body of the whole female reproductive system in locust saline solution side to side with a (I) schematic anatomy of the female reproductive system of another Acrididae, *Locusta migratoria* (adapted from ^28^).

**CRITICAL**: Reproductive tissues may adhere to the fat body on the cuticle; gently detach and reposition any displaced organs.

**4. Expose the reproductive tract.**

**a.** Make a final cut beneath the last abdominal segment connected to the ovipositor and remove the cuticle flap completely (Fig. 14E).
**b.** Gently pull the gut laterally (typically to the left) using fine tweezers, moving it out of the dissection field without puncturing it or cutting it to avoid contamination
**c.** Secure the gut in this position with a pin; the pin should hold the gut back rather than puncturing or piercing it in place (Fig. 14F).
**5. Isolate accessory glands.**

**a.** Remove excess fat body located posterior to the ovarioles using fine tweezers.
**b.** Identify the accessory glands as translucent to whitish paired structures adjacent to the top of the reproductive tract (Fig. 14G).
**c.** Using fine tweezers and small scissors, carefully cut and collect the accessory glands (Fig. 14GHI).
**d.** Transfer immediately into a labeled tube and snap-freeze or place in ice-cold saline as required.
**6. Collect the shared oviducts (if needed).**

**a.** Remove or reposition the pin holding the gut if necessary to improve access.
**b.** Identify the oviducts and their connection to the vagina; exercise caution on the left side where the reproductive tract crosses beneath the gut.
**c.** Cut the V-shaped oviducts above the vagina using fine scissors (Fig. 14HI) and transfer them into a labeled tube for snap-freezing.

### Part 6: Total RNA extraction and validation

Timing: **3h**

This step describes the extraction of high-quality total RNA from locust tissues using magnetic bead–based purification, followed by RNA quantification and integrity assessment. The protocol is optimized for small neural and sensory tissues and relies primarily on an automated extraction kit to ensure rapid, reproducible, and contamination-limited processing. Alternative extraction methods and kits have also been validated in our laboratory and can be provided upon request to the technical lead.

### Clean the bench and prepare RNase-free conditions

**Timing**: 40 min

**1. Decontaminate the work surface.**

a. Wear clean nitrile gloves at all times during bench preparation.
b. Dispense 3% bleach onto the bench using a squeeze bottle, spread evenly with a paper towel, and allow to sit for 1 min.

**CRITICAL:** Avoid splashing bleach onto pipettes or equipment; residual bleach can degrade nucleic acids if not removed.

Rinse thoroughly with distilled water using a squeeze bottle and wipe in circular motions.
Repeat the bleach and rinse steps once more.
Allow the surface to air-dry completely or dry with a clean paper towel.
Finish by wiping the bench with RNaseZap according to the manufacturer’s instructions.

**2. Prepare equipment and reagents.**

a. UV-irradiate pipettes for 30 min.
b. Turn on the Maxwell® RSC instrument.
c. Prepare SimplyRNA™ Tissue Kit cartridges for the number of samples to be processed.
d. Label elution tubes clearly.
e. Prepare lysis solutions according to the manufacturer’s instructions:

i. 1-Thioglycerol/Homogenization Solution and keep on ice.
ii. DNase I solution **Pre-lyse and homogenize tissues Timing**: 30 min (for 16 samples)
**3. Thaw and transfer tissues.**

a. Retrieve snap-frozen screw-cap tubes from −80 °C or liquid nitrogen and thaw on ice.
b. Using wide-bore 1000 µL pipette tips, add 250 µL of chilled 1-thioglycerol/homogenization solution directly to each tube.

**NOTE:** The use of wide-bore or wide-orifice tips from 3b is essential to being able to suspend and suck up tissue without sticking and ensures the use of the entire dissected material.

c. Gently pipette up and down to “fish out” the tissue into the liquid of the tip.
d. Transfer the tissue suspension immediately into a new homogenization tube.

**4. Grind and homogenize tissues.**

a. Choose one homogenization method:

i. MagNA Lyser tube containing green ceramic beads, or
ii. 2 mL round-bottom Safe-Lock tube preloaded with one 5 mm sterile stainless-steel bead.
b. Homogenize using one of the following:

i. MagNA Lyser at 6500 rpm for 30 s, followed by a brief spin-down, or
ii. TissueLyser II at 10 Hz for 30 s, followed by a brief spin-down.
c. Repeat homogenization for an additional 15 s if needed.
d. Place tubes on ice immediately after homogenization.
**5. Lyse and clarify samples.**

a. Add 250 µL of lysis buffer to each homogenized sample.
b. Vortex on medium to medium-high speed for 15 s to mix thoroughly.
c. Spin down briefly to collect the lysate.

### Load samples and run Maxwell RSC extraction

**Timing**: 15 min (for 16 samples)

**6. Transfer lysate to SimplyRNA tissues cartridges.**

a. For MagNA Lyser tubes:
b. Use a pipette tip to carefully mix the lysate.
c. Transfer the supernatant to a SimplyRNA cartridge, ensuring material trapped around beads is recovered.
d. For TissueLyser tubes:
e. Gently mix the lysate by pipetting up and down.
f. Displace the stainless-steel bead slightly and transfer the lysate into the cartridge.
g. Add 10 µL DNase I solution directly into each cartridge well.
**7. Set elution volumes and run RNA extraction.**

a. Adjust elution volume according to tissue type:
b. Optic lobes and fat body: 50 µL
c. Metathoracic ganglia and antennal lobes: adjust to minimal recommended volume
d. Start the SimplyRNA Tissue program on the Maxwell® RSC.

### RNA quantification and quality control

**Timing**: 60 min

**8. Quantify RNA concentration.**

a. Measure RNA concentration using a Qubit fluorometer according to the manufacturer’s instructions, using the High-sensitivity RNA assay.
b. Use 3 µL of eluted RNA per measurement.
**9. Assess RNA integrity.**

a. Analyze a subset of samples using the TapeStation High Sensitivity RNA ScreenTape Analysis according to the manufacturer’s protocol.
b. Interpret profiles with insect-specific considerations.

**NOTE:** When using a 72 °C denaturation step, the typical 18S/28S rRNA ratio will not be visible in insects. Most insect species possess a “hidden break” of the hydrogen bonds in the 28S rRNA, resulting in atypical electropherograms that differ from vertebrate RNA benchmarks ^29^.

**10. Store samples at -80 °C until downstream library preparation.**

### Expected outcomes

When Parts 1 and 2 are followed as described, it is possible to establish and maintain *Schistocerca* colonies under long-term crowded and isolated conditions for multiple consecutive years without the need to regularly introduce new field-collected stock. Under these conditions, colonies show stable reproductive output, predictable development times, and consistent phase-associated phenotypes across generations.

Following Parts 4 and 5, and after sufficient technical training using multiple species, researchers can achieve rapid, reproducible whole-body dissections targeting neural, chemosensory, digestive, and reproductive tissues. With practice, complete dissections of key tissues can be performed in <20 min per individual (Figure 15), while preserving tissue integrity and minimizing handling-induced transcriptional artifacts.

**Figure 15:**
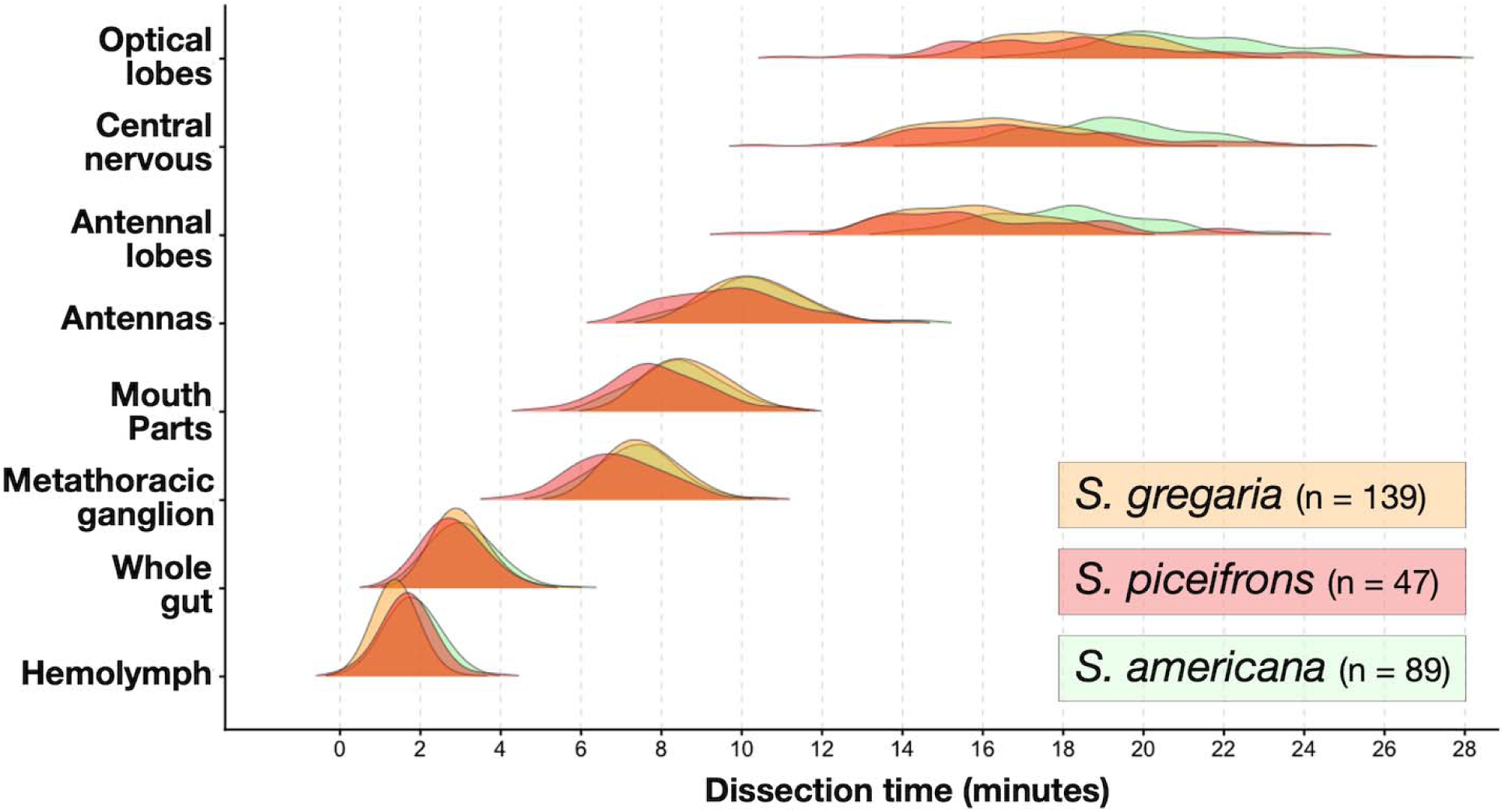
Expected dissections timing distribution for three species of *Schistocerca*. Dissections of whole-body last-instar female nymphs, as described in part 5A, were recorded by one experimenter for S. gregaria and by two for *S. piceifrons* and *S. americana*. While the separation of body segments, metathoracic ganglia, and antennas was similar in timing, the extraction of different central nervous system tissues is a more lengthy process that varies across individuals within the same species and requires extensive practice beforehand.

When tissues are processed live and immediately subjected to magnetic bead–based RNA extraction as described in Part 6, the protocol yields high-quality total RNA suitable for bulk transcriptomic analyses (Figure 16), including from small and fragile tissues such as metathoracic ganglia and antenna lobes. As summarized in Table 2, the amount of total RNA obtained can vary across tissues, with optic lobes and compound eyes providing large amounts, while the lowest amounts are obtained from metathoracic ganglia and antenna lobes. Solitarious females are often larger, and this is reflected by the available RNA quantity in similar tissues compared to gregarious females, especially in *S. gregaria* (Table 2).

**Figure 16:**
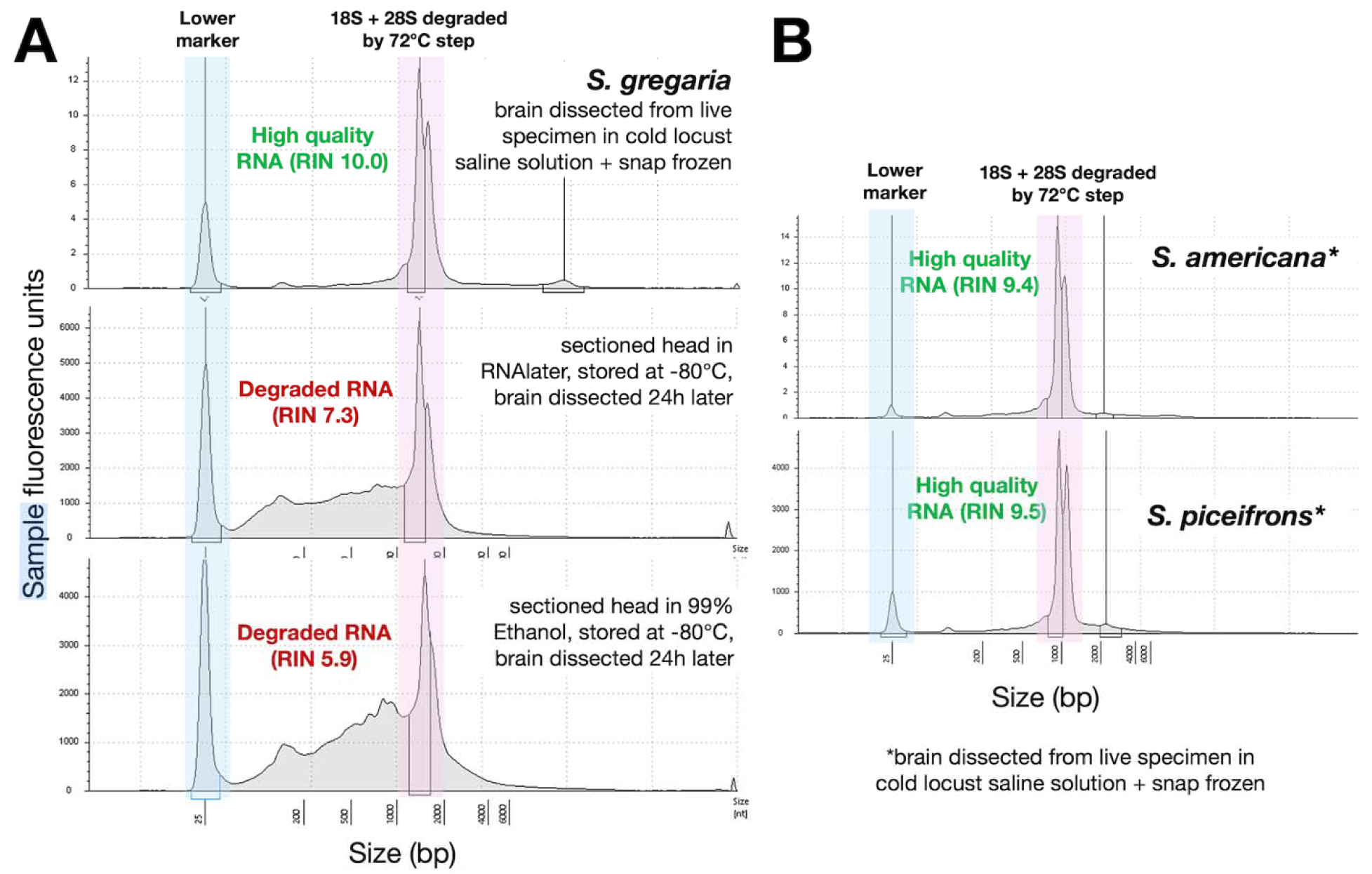
RNA integrity profiles obtained from whole-brain extractions in *Schistocerca* species. (A) Comparison of RNA TapeStation electropherograms from whole brains preserved using RNAlater or >90% ethanol at −80 °C, followed by delayed dissection (24 h). Although convenient, both preservation methods resulted in reduced tissue integrity and partial RNA degradation due to difficulties in accurately isolating the brain. RNAlater caused tissues to become soft and gelatinous, whereas ethanol dehydration produced brittle, flaky tissues, complicating the removal of surrounding structures, including the neural sheath. (B) High-quality RNA profiles obtained from live-dissected brains of *S. gregaria*, *S. americana*, and *S. piceifrons*. Profiles differ from standard vertebrate benchmarks because insect 28S rRNA was degraded and co-migrated with the 18S rRNA peak following 72 °C denaturation during TapeStation analysis.

**Table 2:** Total RNA quantity available from low-input whole tissue extraction using the Maxwell 48 RSC and simplyRNA tissue kit for different species and different density conditions. Values indicate average RNA 土 standard deviation quantified using Qubit HS RNA assays for n specimens.

|  |  |  |  |  |  |
| --- | --- | --- | --- | --- | --- |
|  |  | Metathoracic | Antenna | Mushroom | Optic lobes + |

|  |  | ganglia (*total RNA in ng*) | lobes (*total RNA in ng*) | body and brain (*total RNA in ng*) | compound eyes (*total RNA in µg*) |
| --- | --- | --- | --- | --- | --- |
| *S. gregaria* | Solitarious | 683.7 ± 201.6<br>(n = 59) | 237.5 ± 59.6<br>(n = 59) | 586.2 ± 128.5<br>(n = 53) | 8.98 ± 4.14<br>(n = 54) |
|  | Gregarious | 500.0 ± 233.5<br>(n = 59) | 198.4 ± 93.1<br>(n = 50) | 476.7 ± 176.4<br>(n = 49) | 5.21 ± 1.90<br>(n = 41) |
| *S. americana* | Isolated | 237.8 ± 89.6<br>(n = 60) | 133.7 ± 61.5<br>(n = 40) | 313.9 ± 122.8<br>(n = 54) | 3.13 ± 1.07<br>(n = 60) |
|  | Crowded | 298.9 ± 65.4<br>(n = 60) | 149.4 ± 64.2<br>(n = 59) | 273.1 ± 74.2<br>(n = 60) | 3.32 ± 0.67<br>(n = 60) |
| *S. piceifrons* | Solitarious | 200.3 ± 51.5<br>(n = 30) | 92.7 ± 34.6<br>(n = 16) | 301.1 ± 138.8<br>(n = 30) | 2.22 ± 0.75<br>(n = 30) |
|  | Gregarious | 264.5 ± 80.8<br>(n = 60) | 149.1 ± 57.0<br>(n = 55) | 417.5 ± 86.0<br>(n = 60) | 2.52 ± 0.60<br>(n = 60) |
| *S. serialis cubense* | Isolated | 100.5 ± 28.3<br>(n = 17) | 99.1 ± 42.1<br>(n = 6) | 208.8 ± 90.9<br>(n = 19) | 1.64 ± 0.41<br>(n = 20) |
|  | Crowded | 96.3 ± 25.3<br>(n = 52) | 92.6 ± 27.6<br>(n = 20) | 182.1 ± 104.1<br>(n = 56) | 2.00 ± 0.67<br>(n = 60) |

Using a fraction of these RNA outputs is enough to yield high-quality library preparation using commercial kits, such as the Illumina Stranded Total RNA prep kit with RiboZero depletion.

While such processes and successes are not discussed further in this protocol, ongoing sequencing efforts have been successful but have recognized low RiboZero performance in depleting rRNA from Orthopteran libraries.

### Limitations

**Dissection throughput and temporal constraints:** This protocol requires careful experimental and temporal planning, particularly for time-course experiments that combine density transitions with live dissections. Because tissue integrity and RNA quality depend on rapid processing, dissections must be scheduled to include rest periods for the operator while still maximizing the number of individuals processed per day. Overextended dissection sessions increase the risk of RNA degradation and handling-induced transcriptional artifacts.

Time-course experiments are especially sensitive to unexpected mortality. Individual loss during rearing, transfer, or staging can compromise replicate numbers at specific time points, potentially introducing bias in cohort-based analyses. We therefore strongly recommend staging backup individuals for all time-sensitive experiments. In addition, sluggish or abnormally behaving individuals should be excluded, as these often reflect compromised physiological status and are frequently associated with degraded internal tissues.

**Dependence on live tissue processing:** The workflow relies on live dissection of specimens. Despite extensive testing, preservation of whole individuals in RNAlater or >90% ethanol at −80

°C followed by delayed dissection consistently resulted in reduced tissue integrity and poor RNA quality (Figure 16). RNAlater-treated tissues become soft and gelatinous, whereas ethanol-fixed tissues are brittle and prone to fragmentation, both of which impair accurate separation of neural and sensory tissues. These preservation approaches are therefore not recommended for transcriptomic analyses requiring fine tissue dissection.

**Environmental stability and temperature control:** Environmental stability represents an additional limitation. Although controlled-temperature rooms are sufficient in many settings, rapid external temperature fluctuations (such as those commonly experienced in regions like Texas) can affect building climate control and introduce unintended thermal variation. Climate-controlled chambers provide a more stable and reliable alternative for long-term rearing and experimental standardization.

Importantly, even under constant 32 °C settings, daily photoperiod cycles (12 h light:12 h dark) naturally generate modest day-night temperature differences in the absence of radiant heaters. These fluctuations are acceptable and biologically relevant, as they mimic natural thermal variation experienced by locusts in the field.

**Multi-species rearing, quarantine, and personnel training:** These require rigorous personnel training, careful daily tracking, and strict adherence to labeling and handling protocols.

Inadequate labeling, handling errors, or insufficient awareness of species-specific traits may increase the risk of cross-colony contamination or unintended hybridization. In cases of suspected cross-contamination, affected colonies should be terminated and restarted from verified sources.

In addition, access to quarantine-approved facilities is required for certain *Schistocerca* species and geographic lineages, which may limit adoption of the protocol in some institutions.

Personnel working across multiple species should receive specific training in species identification, reproductive biology, and containment procedures to ensure colony integrity and regulatory compliance.

### Troubleshooting

**Colony health and contamination:** Although the workflow minimizes pathogen transmission through cage design and routine cleaning, feeding and handling (especially in crowded colonies) create potential contamination pathways via contact with feces and external surfaces. Early signs of disease (e.g., reduced feeding, lethargy, abnormal coloration) should be addressed immediately. Non-organic commercially bought food may contain pesticide which can cause immediate colony-wide deaths with symptoms such as twitching.

In cases of suspected disease outbreaks (Figure 17), opportunistic entomopathogens *Pseudomonas aeruginosa* and *Pseudomonas sp.* ^30^ were found abundant in decaying locust carcasses (Peterson, unpublished), and treatment or cage termination measures may be necessary. Spraying food sources daily for 5-7 days with a triple-sulfate-based solution can reduce the occurrence of gram-negative bacteria and isolate affected cages while monitoring unaffected colonies. When recurrent or unexplained symptoms occur, molecular identification of pathogens is strongly advised before colony-wide intervention.

**Figure 17:**
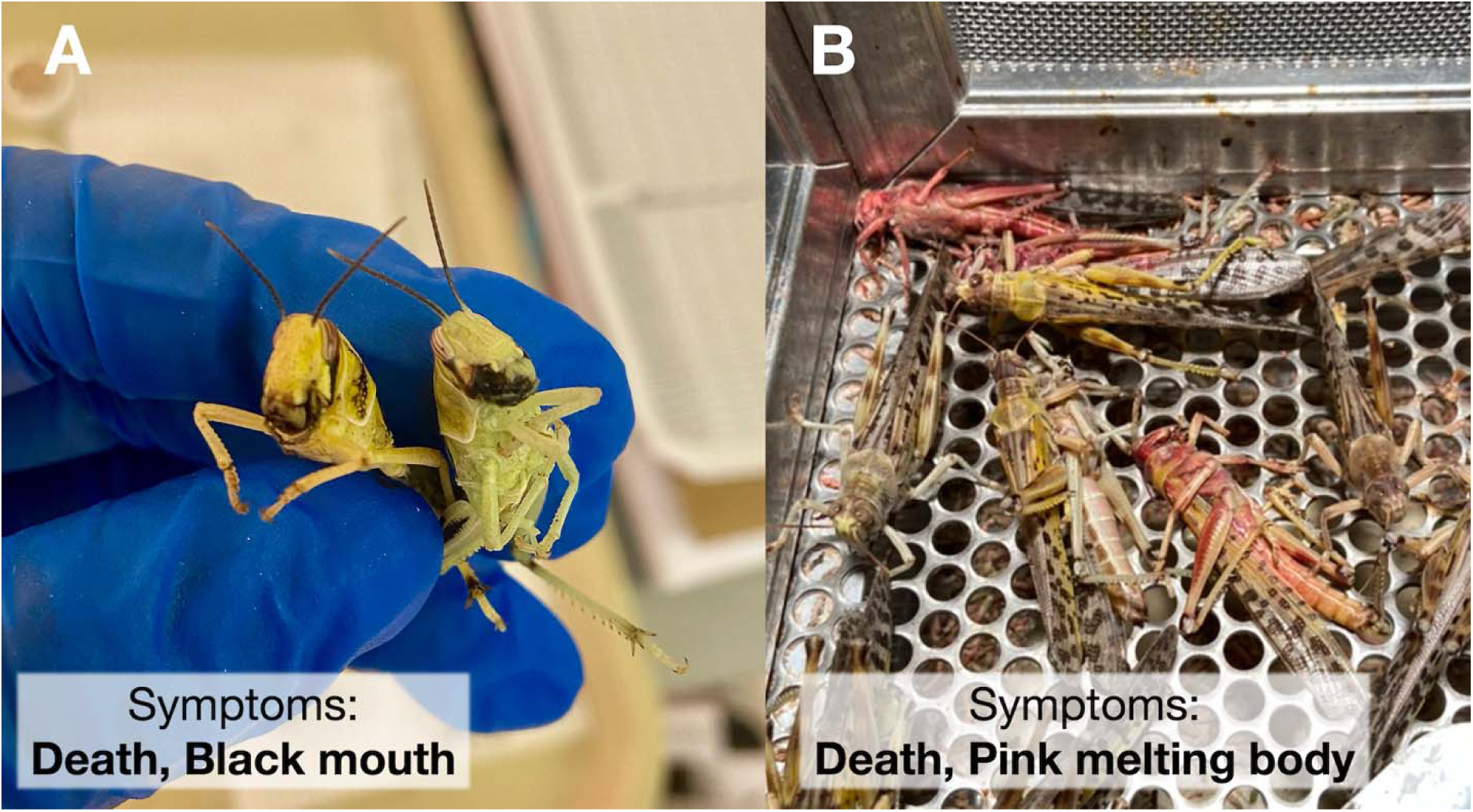
Examples of colony disease and bacterial outbreaks that can occur via food infection and enhanced by cannibalism of dying insects with symptoms such as (A) necrosis of the digestive system and (B) body degradation leading to a pink color and soft tissues.

**Time-course dissection bottlenecks:** If dissections fall behind schedule, reduce the number of tissues collected per individual rather than extending handling time. Prolonged dissections increase the risk of RNA degradation and reduce reproducibility. Neural sheath removal, eye detachment, and gut separation vary across species and phase states. For example, our experience revealed that dissecting isolated *S. gregaria* is easier than dissecting crowded ones, due to differences in size and internal structure. In parallel, the neural sheath is more difficult to remove in *S. piceifrons* than in *S. gregaria*. Training should include cross-species testing before initiating large-scale experiments to identify problematic steps.

## Author contributions

MAT collected data, designed and optimized protocols for marking, time-course dissections of nymph insects, molecular processing and sequencing, produced figures, and wrote the manuscript. VAPS collected data, designed and optimized protocols for time-course dissections of adult insects, produced figures, and wrote the manuscript. BW generated high-quality pictures of insect development and wrote the manuscript. RM collected data, optimized rearing protocols, managed crowded and isolated insects, and wrote the manuscript. CB optimized gut dissection protocols and wrote the manuscript. AMCM optimized behavioral assay protocols and wrote the manuscript. JBL tested and optimized RNA extraction for subsequent molecular processing and wrote the manuscript. STB designed and managed insect colonies, optimized gut dissection protocols, acquired funding, and wrote the manuscript. GAS designed and managed insect colonies, optimized behavioral assays, acquired funding, and wrote the manuscript. HS designed and managed insect colonies, designed dissection and time course experiments, acquired fundin,g and wrote the manuscript.

## Resource availability

### Lead contact

Hojun Song

### Technical contact

Maeva Techer

### Materials availability

*Schistocerca* colonies are maintained in quarantine facilities but can be visited on-site at Texas A&M University upon contact with the lead contact, GAS, and STB.

### Data and code availability

All data used for this protocol have been made available directly in the manuscript. Individual high-resolution images of the daily development of *Schistocerca* locusts and grasshoppers are available upon contact with the technical contact and BW.

## Acknowledgments

This research is a major product of the Behavioral Plasticity Research Institute (BPRI), which was funded by the National Science Foundation (NSF)’s Biology Integration Institutes (BII) program (DBI-2021795 to HS and STB). Additional funding was provided by the NSF IOS-1253493 to HS. Importation of locusts was permitted by USDA APHIS PPQ permits (P526P-15-03851, P526P-19-02151, and P526P-21-06855 to HS). We thank Arianne Cease and Rick Overson at Arizona State University for providing seed colonies of *S. gregaria*. We thank the students who actively participated in improving and daily testing the colony care and time-experiment setup: Alyssa Canova, Serena Farell, Tessa Huddleston, Kate Wilson, Corinne Allen, Reese Roberson, Kendall Walton, and Katie Puperi. We extend our warm thanks to former colony manager Helen Vasquez, who helped maintain and produce isolated lines for the data acquired. Finally, we thank Fabrizio Gabbiani and Richard Dewell at Baylor College of Medicine for providing guidance and initial training in central nervous system dissections of *S. americana*.

## Declaration on interests

The authors declare no conflict of interest. The funders had no role in the design of the study, the collection, analysis, or interpretation of the data, or the writing of the manuscripts.

## Declaration of generative AI and AI-assisted technologies in the manuscript preparation process

During the preparation of this work, the authors used Grammarly and ChatGPT5.2 solely to assist with English grammar and language editing. After using these tools, the authors carefully reviewed and edited the content as needed and take full responsibility for the content of the published article. <u>Illustrate.co</u> was used to transform the authors’ original photographic images into textbook-style, realistic scientific illustrations.

## Notes

### Competing Interest Statement

The authors have declared no competing interest.

## References

1. Simpson, S.J., and Sword, G.A. (2008). Locusts. Curr. Biol. 18, R364–R366.

2. Foquet, B., Little, D.W., Medina-Durán, J.H., and Song, H. (2022). The time course of behavioural phase change in the Central American locust *Schistocerca piceifrons*. J. Exp. Biol. 225, jeb244621.

3. Rogers, S.M., Cullen, D.A., Anstey, M.L., Burrows, M., Despland, E., Dodgson, T., Matheson, T., Ott, S.R., Stettin, K., Sword, G.A., et al. (2014). Rapid behavioural gregarization in the desert locust, *Schistocerca gregaria* entails synchronous changes in both activity and attraction to conspecifics. J. Insect Physiol. 65, 9–26.

4. Pflüger, H.-J., and Bräunig, P. (2021). One hundred years of phase polymorphism research in locusts. J. Comp. Physiol. A Neuroethol. Sens. Neural Behav. Physiol. 207, 321–326.

5. Latchininsky, A.V., Sergeev, M.V., and Fedotova, A.A. (2021). The centennial of Sir Boris Uvarov’s locust phase theory. Metaleptea.

6. Guo, X., and Kang, L. (2024). Phenotypic plasticity in locusts: Trade-off between migration and reproduction. Annu. Rev. Entomol. 10.1146/annurev-ento-013124-124333.

7. Wang, X., and Kang, L. (2014). Molecular mechanisms of phase change in locusts. Annu. Rev. Entomol. 59, 225–244.

8. Simpson, S.J. (2022). A journey towards an integrated understanding of behavioural phase change in locusts. J Insect Physiol 138, 104370.

9. Cullen, D.A., Cease, A.J., Latchininsky, A.V., Ayali, A., Berry, K., Buhl, J., De Keyser, R., Foquet, B., Hadrich, J.C., Matheson, T., et al. (2017). Chapter Seven - From Molecules to Management: Mechanisms and Consequences of Locust Phase Polyphenism. In Advances in Insect Physiology, H. Verlinden, ed. (Academic Press), pp. 167–285.

10. Hinks, C.F., and Erlandson, M.A. (1994). Rearing Grasshoppers and Locusts: Review, Rationale and Update. Journal of Orthoptera Research, 1–10.

11. Berthier, K., Chapuis, M.-P., Simpson, S.J., Ferenz, H.-J., Kane, C.M.H., Kang, L., Lange, A., Ott, S.R., Ebbe, M.A.B., Rodenburg, K.W., et al. (2010). Chapter 1 - Laboratory Populations as a Resource for Understanding the Relationship Between Genotypes and Phenotypes: A Global Case Study in Locusts. In Advances in Insect Physiology, S. J. Simpson, ed. (Academic Press), pp. 1–37.

12. Pfennig, D.W. (2021). Phenotypic plasticity & evolution: Causes, consequences, controversies D. W. Pfennig, ed. (CRC Press).

13. Simpson, S.J. & Sword, G.A. (2009). Phase polyphenism in locusts: Mechanisms, population consequences, adaptive significance and evolution. In Phenotypic Plasticity of Insects: Mechanisms and Consequences. (Science Publishers Inc., Plymouth), pp. 147–190. 10.1201/b10201.

14. Simpson, S.J., Sword, G.A., and Lo, N. (2011). Polyphenism in insects. Curr. Biol. 21, R738–R749.

15. Corona, M., Libbrecht, R., and Wheeler, D.E. (2016). Molecular mechanisms of phenotypic plasticity in social insects. Curr Opin Insect Sci 13, 55–60.

16. Uvarov, B.P. (1921). A revision of the genus Locusta, L. (= Pachytylus, Fieb.), with a new theory as to the periodicity and migrations of locusts. Bull. Entomol. Res. 12, 135–163.

17. Anstey, M.L., Rogers, S.M., Ott, S.R., Burrows, M., and Simpson, S.J. (2009). Serotonin mediates behavioral gregarization underlying swarm formation in desert locusts. Science 323, 627–630.

18. Song, H., Foquet, B., Mariño-Pérez, R., and Woller, D.A. (2017). Phylogeny of locusts and grasshoppers reveals complex evolution of density-dependent phenotypic plasticity. Sci. Rep. 7, 6606.

19. Foquet, B., Castellanos, A.A., and Song, H. (2021). Comparative analysis of phenotypic plasticity sheds light on the evolution and molecular underpinnings of locust phase polyphenism. Sci. Rep. 11, 11925.

20. Peng, W., Ma, N.L., Zhang, D., Zhou, Q., Yue, X., Khoo, S.C., Yang, H., Guan, R., Chen, H., Zhang, X., et al. (2020). A review of historical and recent locust outbreaks: Links to global warming, food security and mitigation strategies. Environ. Res. 191, 110046.

21. Le Gall, M., Overson, R., and Cease, A. (2019). A global review on locusts (Orthoptera: Acrididae) and their interactions with livestock grazing practices. Front. Ecol. Evol. 7. 10.3389/fevo.2019.00263.

22. Nishide, Y., Tanaka, S., and Saeki, S. (2015). Egg hatching of two locusts, *Schistocerca gregaria* and *Locusta migratoria*, in response to light and temperature cycles. J. Insect Physiol. 76, 24–29.

23. Shand, J.D. (2016). The effects of density-dependent polyphenism on circadian biology of the desert locust *Schistocerca gregaria*.

24. Dong, Y., and Friedrich, M. (2005). Nymphal RNAi: systemic RNAi mediated gene knockdown in juvenile grasshopper. BMC Biotechnol. 5, 25.

25. Roonwal, M.L., and Imms, A.D. (1997). Variation and structure of the eyes in the desert locust, Schistocerca gregaria (Forskål). Proceedings of the Royal Society of London. Series B - Biological Sciences 134, 245–272.

26. Castillo, A.E., Rossoni, S., and Niven, J.E. (2018). Matched Short-Term Depression and Recovery Encodes Interspike Interval at a Central Synapse. Sci. Rep. 8, 13629.

27. Staudacher, E.M., Cigan, M.-L., Wenz, F., Pollun, A., Beck, S., Beck, M., Reh, F., Haas, J., and Homberg, U. (2023). Organization of descending neurons in the brain of the desert locust. J. Comp. Neurol. 531, 1350–1380.

28. Lange, A.B. (2009). The female reproductive system and control of oviposition in *Locusta migratoria migratorioides*. Can. J. Zool. 87, 649–661.

29. Winnebeck, E.C., Millar, C.D., and Warman, G.R. (2010). Why does insect RNA look degraded? J. Insect Sci. 10, 159.

30. Ashrafi, S.H., Zuberi, R.I., and Hafiz, S. (1965). Occurrence of *Pseudomonas aeruginosa* (Schroeter) Migula as a pathogenic bacterium of the desert locust, *Schistocerca gregaria* (Forskål). J. Invertebr. Pathol. 7, 189–191.

